# An expanded palette of bright and photostable organellar Ca^2+^ sensors

**DOI:** 10.1101/2025.01.10.632364

**Authors:** Agathe Moret, Helen Farrants, Ruolin Fan, Kelsey G. Zingg, Bryon Silva, Camilla Roselli, Thomas G. Oertner, Christine E. Gee, Dafni Hadjieconomou, Vidhya Rangaraju, Eric R. Schreiter, Jaime de Juan-Sanz

## Abstract

The use of fluorescent sensors for functional imaging has revolutionized the study of organellar Ca^2+^ signaling. However, understanding the dynamic interplay between intracellular Ca^2+^ sinks and sources has been hindered by the lack of bright, photostable, and multiplexed measurements in different organelles, limiting our ability to define how Ca^2+^ shapes cell physiology across fields of biology. Here we introduce a new toolkit of chemigenetic organellar Ca^2+^ indicators whose color is tunable by reconstituting their fluorescence with different exogenous rhodamine dye-ligands, which significantly expand the capacity for multiplexing organellar Ca^2+^ measurements. These sensors, which we named ER-HaloCaMP and Mito-HaloCaMP, are optimized to report Ca^2+^ dynamics in the endoplasmic reticulum (ER) and mitochondria of mammalian cells and neurons, and show significantly improved brightness, photostability and responsiveness when compared to current best-in-class alternatives. Using either red or far-red dye-ligands, both ER-HaloCaMP and Mito-HaloCaMP enable visualizing ER and mitochondrial Ca^2+^ dynamics in neuronal axons, a subcellular location that only contains a few ER tubules and small mitochondria, structural limitations that have impaired measurements with previous red sensors. To show the expanded multiplexing capacities of our toolkit, we measured interorganellar Ca^2+^ fluxes simultaneously in three different subcellular compartments in live cells, revealing that the amplitude of ER Ca^2+^ release controls the efficacy of ER-mitochondria Ca^2+^ coupling in a cooperative manner. Organellar HaloCaMPs enable also measuring Ca^2+^ dynamics in intact brain tissue from flies and rodents, demonstrating their versatility across biological models. Our new toolkit provides an expanded palette of bright, photostable and responsive organellar Ca^2+^ sensors, which will facilitate future studies of intracellular Ca^2+^ signaling across fields of biology in health and disease.

## Introduction

The ability of cells to generate transient Ca^2+^ fluctuations in subcellular locales is fundamental for controlling essential cellular processes such as gene expression, cell metabolism or neuronal communication^1,2^ Organelles such as mitochondria or the ER can function within cells as sinks or sources of Ca^2+^. They modulate Ca^2+^ dynamics across cellular compartments with temporal and spatial precision, shaping cell physiology^3,4^. For ample, Ca^2+^ transfer between the ER and mitochondria accelerates mitochondrial metabolism to fuel energy-intensive tasks^5^, modulates mitochondrial fission-fusion dynamics^6^,7 or influences central cellular processes such as autophagy and stress responses^8^,^9^ On the other hand, mitochondria and ER uptake Ca^2+^ in neurons during neurotransmission, shaping the capacity of neuronal communication^10-14^. Not surprisingly, owing to the critical role of ER and mitochondrial Ca^2+^ signaling in both physiological and pathological processes^3, 15-17^, their investigation has garnered significant attetinon over the past several decades^2 18^.

For many years, insights into organellar Ca^2+^ functions were primarily obtained using indirect cytosolic Ca^2+^ measurements coupled with organelle manipulations’. However, recent advancements now enable measuring Ca^2+^ dynamics directly within the ER or mitochondria using adapted genetically encoded sensors (GECis)<sup>^11, 12 19-22^Among previous mitochondria and ER Ca^2+^sensors, GCaMP variants have stood out due to their high dynamic range and sensitivity^11-14^ -^19-23^ -^24^ However, for intraorganellar use they have inherent limitations. First, their excitation spectrum overlaps with other commonly used fluorescent proteins, sensors and optogenetic tools2s--^21^, complicating multicolor imaging needed for dissecting complex cellular processes; second, blue light excites autofluorescence in biological tissues^28^, making it difficult to detect calcium dynamics in small organellar structures with green fluorescent indicators^12 22^ third, imaging with blue light is associated with higher phototoxicity^29^,^30^ which may complicate measurements over time. These limitations suggest that red-shifted GECls could provide a reliable imaging alternative for imaging Ca^2+^dynamics in organelles. However, despite recent advances, available red-shifted GECls for ER and mitochondrial Ca^2+^ imaging are not ideal due to either an inappropriately high Ca^2+^ affinity2^1^ or a limited dynamic range^19^,^20^ Additionally, the lack of far-red Ca^2+^sensors for these organelles has further limited simultaneous multi-wavelength imaging of organellar and cytosolic Ca^2+^ dynamics, along with other key physiological markers, limiting multi-compartment dissections of Ca^2+^ fluxes and their relevance for cellular physiology in health and disease.

Leveraging HaloTag^31^ as a scaffold for generating novel sensors has provided several novel indicators that enable measuring diverse pieces of biology in different colors, as the color of emission is provided by different membrane-permeable rhodamine dye-ligands that irreversibly bind to the HaloTag. These include sensors for voltage^32,33^, ATP34, NAD+, ^34^, dopamine^35^, pH^36^ and protein aggregation^37^, among others^38,39^. However, while several HaloTag-based strategies have also provided robust multicolor cytosolic Ca^2+^ sensors40-4^3^, suitable versions to expand the palette of organellar Ca^2+^sensors do not exist. In this work, we present a new suite of chemigenetic indicators derived from HaloCaMP, a genetically encoded Ca^2+^ sensor whose fluorescence is provided by membrane-permeable Janelia Fluor rhodamine dye-ligands with a choice of colors appended to the HaloTag Ligand (JF-HTL)^40^. We generated novel organellar HaloCaMP sensors, tailored specifically for Ca^2+^ measurements in ER and mitochondria: ER-HaloCaMP and Mito-HaloCaMP. These sensors combine robust responsiveness with the enhanced brightness and photostability provided by JF-HTLs. Along with these advantages, the indicators feature tunable red and far-red emission, enabling multicolor imaging without the spectral overlap or interference issues commonly associated with green GECis. We compared ER-HaloCaMP and Mito-HaloCaMP to existing red-emitting organellar sensors, and found they responded significantly better (1.6-to 2.3-fold) and were significantly brighter (10-to 50-fold). Physiological responses of ER-HaloCaMP reconstituted with JF_585_-HTL (ER-HaloCaMP_585_) in cells and neurons were indistinguishable from the those of the gold-standard green ER Ca^2+^ sensor ER-GCaMP6-210^1^2, but the photostability and brightness-over-background of ER-HaloCaMP_585_ were significantly improved, making it the ideal sensor for ER Ca^2+^ measurements. This work provides a new technological framework to enable novel investigations on cellular Ca^2+^ fluxes, offering new insights into the fundamental role of Ca^2+^ signaling in health and disease.

## Results

### Developing a bright and highly responsive red ER Ca^2+^ sensor based on HaloCaMP

The ER contains ∼2000 times more Ca^2+^ than the cytosol. This surplus of Ca^2+^ inevitably saturates all existing red cytosolic Ca^2+^ GECis ^44-49^ making them unresponsive to changes in ER Ca^2+^. To overcome this issue and image ER Ca^2+^ in red, previous efforts have screened for mutations that lower the affinity of red GECI to match resting ER Ca^2+^ levels. These approaches generated LAR-GECOs, which preserved a good dynamic range yet their affinity for Ca^2+^ was not lowered enough to match ER Ca^2+^ levels, limiting their usability2^1^. Alternative efforts generated R-CEPIAer, which presented low affinity but its responsiveness remained ten times lower compared to that of green ER Ca^2+^ GECls^19^.

To overcome the limitations of existing red ER Ca^2+^ sensors and expand their spectral range beyond red, we first generated a series of HaloCaMP1a variants harboring mutations in key Ca^2+^-coordinating sites to lower their affinity (Fig. lA; Supp. Fig. SlA; Table 1). We then performed Ca^2+^ titrations and identified a variant that presented both high dynamic range (∼15 ΔF/F_0_) and low affinity (EC_50_ = 86 µM), which we named Low Affinity (LA)-HaloCaMP_585_ (Fig. 1B). The general properties of this low affinity variant did not significantly change when Ca^2+^ titrations were performed at 37 °C, supporting its usability for live cell experiments (Supp. Fig. SlC; EC_50_ = 89 µM, ∼17 ΔF/F_0_). As a control, we confirmed that micromolar/millimolar increases in Ca^2+^ did not cause any increase in fluorescence of HaloTag reconstituted by JF_585_-HTL (HaloTa&_85_; Fig. 1B). Using purified LA-HaloCaMP_585_, we confirmed that its excitation and emission spectra yielded a peak excitation at 593 nm and emission at 607 nm, confirming its usability as a red sensor (Supp. Fig. SlD).

**Table 1.**
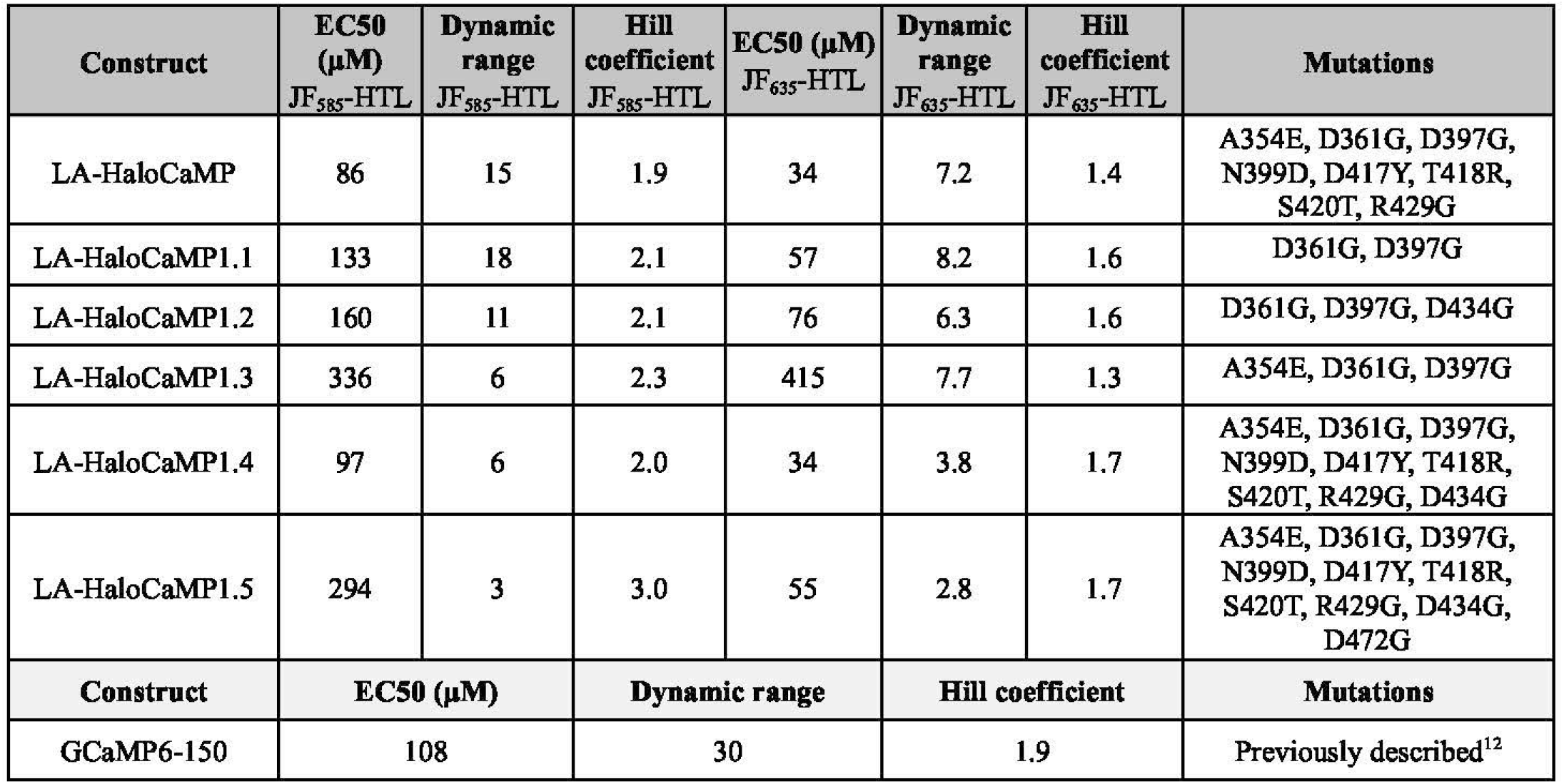
Properties of low affinity variants measured in purified protein at room temperature.

**Table 2.** Biophysical properties of purified LA-HaloCaMP in comparison to previous sensors.

| LA-HaloCaMP | Abs<sub>apo</sub> (nm) | Abs<sub>sat</sub> (nm) | Ex/Em<sub>apo</sub> (nm) | Ex/Em<sub>sat</sub> (nm) | ε<sub>apo</sub> (x1000) (M<sup>-1</sup> cm<sup>-1</sup>) | ε<sub>sat</sub> (x1000) (M<sup>-1</sup> cm<sup>-1</sup>) | Φ<sub>apo</sub> | Φ<sub>sat</sub> | Brightness Apo (mM<sup>-1</sup> cm<sup>-1</sup>) | Brightness Sat (mM<sup>-1</sup> cm<sup>-1</sup>) |
| --- | --- | --- | --- | --- | --- | --- | --- | --- | --- | --- |
| JF<sub>585</sub>-HTL | 591 | 592 | 593/609 | 593/607 | 8 | 129 | N.D. | 0.70 | N.D. | 90 |
| JF<sub>635</sub>-HTL | 642 | 641 | 643/655 | 641/654 | 15 | 74 | N.D. | 0.65 | N.D. | 48 |
| GECIs | Abs<sub>apo</sub> (nm) | Abs<sub>sat</sub> (nm) | Ex/Em<sub>apo</sub> (nm) | Ex/Em<sub>sat</sub> (nm) | ε<sub>apo</sub> (x1000) (M<sup>-1</sup> cm<sup>-1</sup>) | ε<sub>sat</sub> (x1000) (M<sup>-1</sup> cm<sup>-1</sup>) | Φ<sub>apo</sub> | Φ<sub>sat</sub> | Brightness Apo (mM<sup>-1</sup> cm<sup>-1</sup>) | Brightness Sat (mM<sup>-1</sup> cm<sup>-1</sup>) |
| LAR-GECO1 | 574 | 561 | 598 | 589 | 5 | 36 | 0.13 | 0.2 | 0.7 | 7.2 |
| R-CEPIA1<sub>er</sub> | 576 | 562 | 593 | 584 | 5 | 35 | 0.08 | 0.18 | 0.5 | 6.2 |

N.D. = not determined. The photon output of the sensor was too low to accurately determine quantum yield in the HTL = HaloTag ligand. apo = in presence of NTA. sat = in presence of saturating Ca<sup>2+</sup>. Ex = excitation maximum. Em = emission maximum. ε = molar extinction coefficient. Φ = quantum yield. Brightness (mM<sup>-1</sup> cm<sup>-1</sup>) = the product of molar extinction coefficient × quantum yield divided by 1000.

**Figure 1.**
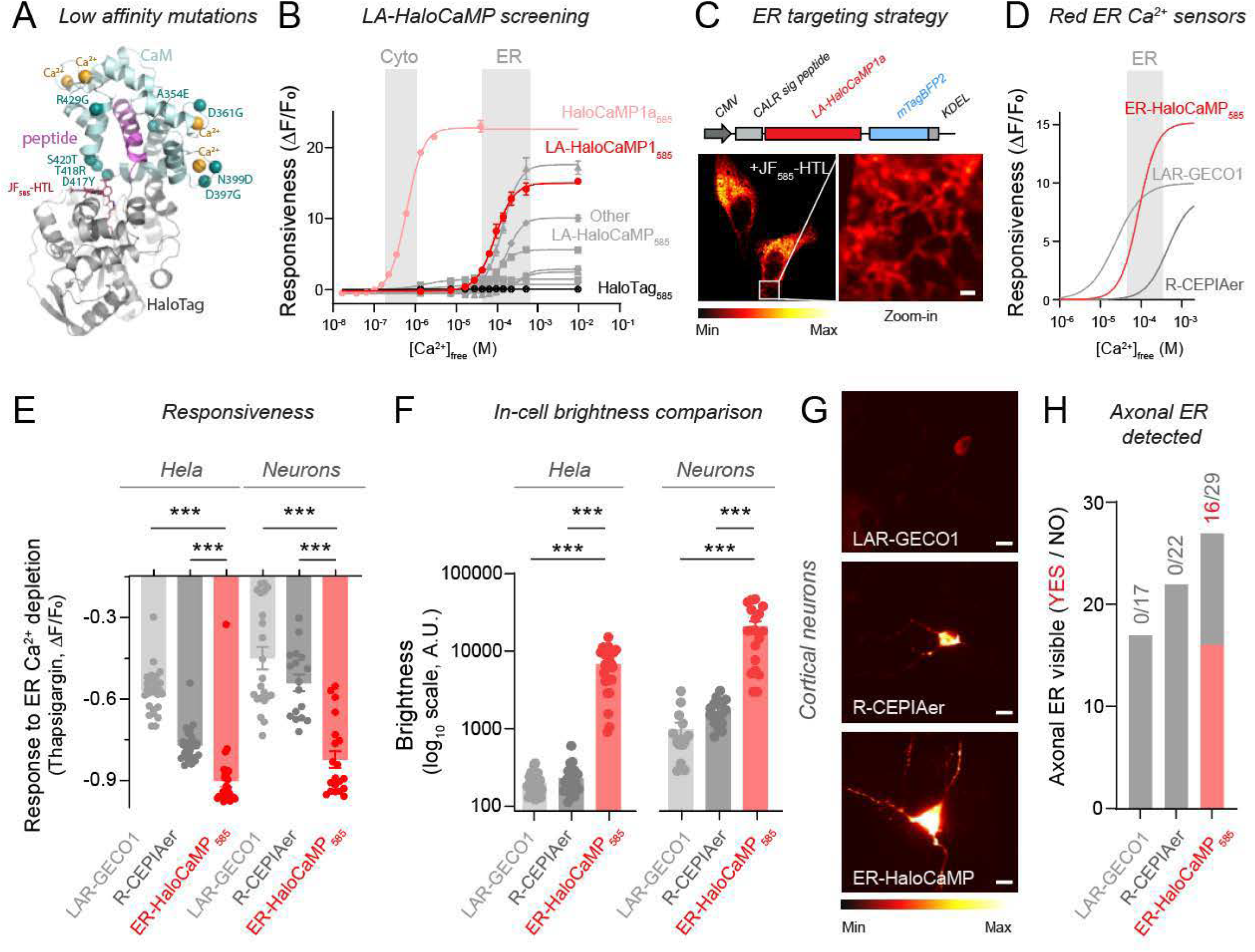
ER-HaloCaMP_585_ is a bright and highly responsive red ER Ca^2+^ sensor. (A) Chai Discovery-predicted structure of low affinity HaloCaMP showing the mutations in the CaM domain in turquoise, Ca^2+^ binding sites in yellow and JF_585_*-HTL* in red. The HaloTag is shown in grey. **(B)** Screening for low affinity HaloCaMP variants: *in vitro* Ca^2+^ titrations of different purified HaloCaMP variants conjugated with *JF*_585_*-HTL*. (C) Top: targeting scheme for expression in the ER by adding the N-terminal signal peptide of calreticulin (CALR sig peptide) and the C-terminal KDEL retention motif. Bottom: high resolution image of a HeLa cell expressing ER-localized LA-HaloCaMP reconstituted with *JF*_585_*-HTL* shows the ER structure. Pseudocolor scale shows low to high fluorescence intensity. Scale bar, 1 µm. **(D)** Comparative responsiveness ofER-HaloCaMP_585_, LAR-GECOl and R-CEPIAer. (E) Quantification ofthapsigargin response (1 µM) in both HeLa cells and cortical neuron somas. **(F-H)** Comparison of relative brightness of LAR-GECO, R-CEPIAer and ER-HaloCaMP_585_ using identical illumination and detection conditions in either HeLa cells or cortical neurons. (F) Quantification of sensor brightness in intact HeLa cells or cortical neuron somata. (G) Representative images of brightness level in neuronal somata. Scale bar, 10 µm (H) Quantificationof the ability to detect ER tubules in axons using red ER calcium sensors. Data are represented as mean ± SEM. See Supplementary Table STl for details on statistical tests and sample sizes.

We next targeted LA-HaloCaMP to the lumen of the ER by flanking it with a modified calreticulin signal peptide and a KDEL retention motif, as successfully used for ER-GCaMP6^12^. This construct, which also was fused to mTagBFP2, was efficiently expressed in the ER of HeLa cells. We confirmed that JF_585_-HTL successfully traversed both the plasma and ER membranes, demonstrating that conjugation between sensor and dye-ligand in the ER is possible (Fig. 1C). We performed *in-cell* Ca^2+^ titrations in permeabilized HEK cells and found that ER-HaloCaMP presents an affinity of 115 µMin cells (Supp. Fig SlD), slightly lower than purified protein. Such a shift between purified and *in-cell* sensor properties is expected and has been observed for ER-GCaMP6 variants^50^. Given the higher dynamic range of ER-HaloCaMP and its better suited affinity for ER Ca^2+^ than LAR-GECO and R-CEPIAer in purified protein (Fig. 1D), we next sought to compare its responsiveness against these sensors under common ER Ca^2+^ perturbations, such as thapsigargin-induced ER Ca^2+^ depletion^51^. Quantifying responses in both HeLa cells and somata of primary cortical neurons, we observed that although LAR-GECO and R-CEPIAer responded to thapsigargin as previously reported^19,21,52^, ER-HaloCaMP_585_ exhibited a significantly larger response in both cell types (Fig. 1E).

We next compared the brightness of ER-HaloCaMP_585_ against previous red ER Ca^2+^ sensors. While the ER is abundant and easily detectable when measured in bulk within a cell, the small ∼60 nm diameter of individual ER tubules can hinder ER Ca^2+^ detection in subcellular locales. This problem becomes apparent when studying neuronal projections, as only a single narrow ER tubule may be present in axons^53,54^ or dendritic spines^54^,^55^ We thus reasoned that sensors with increased brightness should facilitate studying ER Ca^2+^ in subcellular compartments in neurons and other cells. We first compared the molecular brightness in the Ca^2+^-saturated state of ER-HaloCaMP_585_ against previous red ER Ca^2+^ sensors. Purified ER-HaloCaMP_585_ reconstituted with JF_585_-HTL exhibited a brightness of 90 mM-^1^cm-1, substantially higher than LAR-GECOl (7.2 mM-^1^cm-^1^) and R-CEPIAer (6.2 mM-^1^cm-^1^) (Table 1), confirming a nearly 15-fold increase in molecular brightness relative to currently available red GECls. We next expressed LAR-GECO, R-CEPIAer and ER-HaloCaMP_585_ in HeLa cells and primary cortical neurons and using identical illumination and detection conditions we compared their relative *in-cell* brightness. We found that ER-HaloCaMP_585_ was approximately 100-fold brighter in HeLa cells and 20-fold brighter in somata of primary neurons (Fig. 1F, G), supporting data from purified protein. Importantly, these differences in live cell imaging cannot be ascribed to our optical setup, as it is actually better matched to excite and collect fluorescence from LAR-GECOl and R-CEPIAer (see Methods). This suggests that the enhanced detectability of ER-HaloCaMP_585_ in cells is driven by its intrinsically higher brightness.

Next, we investigated the ability of these sensors to detect ER fluorescence within the single, narrow ER tubules of axons, a task that represents a significant challenge in ER Ca^2+^ imaging. This comparison provided a substantial improvement over previous technologies, as neither LAR-GECO nor R-CEPIAer enabled axonal ER detection in any neuron presenting somatic ER labeling, whereas ER-HaloCaMP_585_ allowed us to identify it clearly in over 50% of the transfected neurons when using identical imaging conditions (Fig. 1G, H; see Methods). These results indicate that ER-HaloCaMP_585_ provides a significant improvement in red ER Ca^2+^imaging by providing significantly better responsiveness and brightness.

### ER-HaloCaMP enables ER Ca^2+^ measurements in locales with low ER content

As ER-HaloCaMP_585_ is the only red ER Ca^2+^sensor visible in axons of neurons, we sought to evaluate its responsiveness in this small compartment. When neurons fire action potentials, axonal ER uptakes Ca^2+^ from the cytosol through sarcoplasmic/endoplasmic reticulum Ca^2+^-ATPases (SERCAs)^10, 12^, ^56^ Using field stimulation (see Methods), we electrically stimulated primary cortical neurons to mimic physiological firing paradigms of neurons *in vivo*, such as firing at 20 Hz for ls^57^, and quantified axonal ER Ca^2+^ dynamics using ER-HaloCaMP_585_. We observed robust activity-driven ER Ca^2+^ uptake, which was completely silenced when inhibiting SERCA pumps using thapsigargin, confirming the specificity of the measurements (Fig. 2A, B). Since other red ER Ca^2+^ sensors were not visible in axons, we compared the performance of ER-HaloCaMP_585_ with the current gold standard ER Ca^2+^ sensor, the GFP-based ER-GCaMP6-210^12^. ER-HaloCaMP_585_ exhibited uptake responses comparable to those of ER-GCaMP6-210, demonstrating its high efficacy in measuring ER Ca^2+^ dynamics within small ER volumes (Fig. 2C, D).

**Figure 2.**
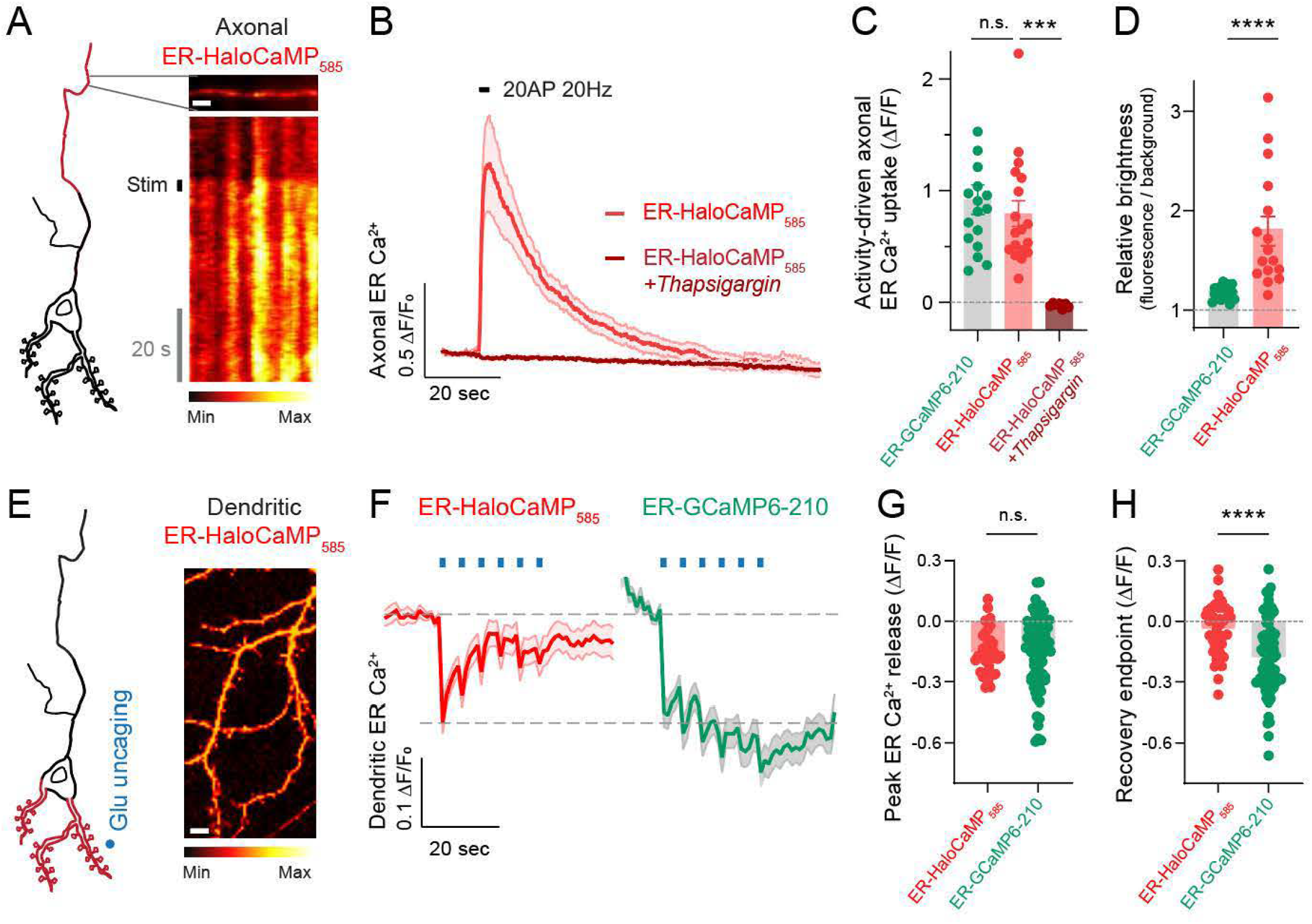
ER-HaloCaMP_585_ enables ER Ca^2+^ measurements in subcellular locales with low ER content. **(A-D)** Axonal ER Ca^2+^ responses to stimulation of 20 action potentials (AP) at 20 Hz measured using different indicators. (A) On the left: schematic representation ofER-HaloCaMP_585_ expressed within a single ER tubule of axon. Right panel shows a representative image of an axon expressing ER-HaloCaMP_585_ and changes in fluorescence induced by stimulation. Scale bar, 4 µm. Pseudocolor scale of intensity shown below. **(B)** Axonal ER Ca^2+^ responses to stimulation of 20AP at 20 Hz before and after 15 minute thapsigargin treatment using ER-HaloCaMP_585_. (C) Quantification of activity-driven ER Ca^2+^uptake peak response upon 20 AP (20 Hz) stimulation using ER-GCaMP6-210 (green), ER-HaloCaMP_585_ (red) and ER-HaloCaMP_585_ (dark red) in the presence of thapsigargin. **(D)** Quantification of relative brightness measured with ER-GCaMP-210 or ER-HaloCaMP_585_ in axons. **(E-H)** Dendritic ER Ca^2+^responses to glutamate uncaging in spines. (E) On the left: schematic representation of ER-HaloCaMP_585_ in dendrites and location of glutamate uncaging on a spine. Right panel shows the dendrites of a neuron expressing ER-HaloCaMP_585_, (pseudocolor scale relative to the image showing low to high intensity). Scale bar, 4 µm. **(F)** Dendritic ER Ca^2+^ responses adjacent to the site of glutamate uncaging measured with ER-HaloCaMP_585_ or ER-GCaMP-210. Blue ticks indicate the 6 uncaging pulses of 100 ms at 0.25 Hz, 7.2 mW 720 nm uncaging pulses. **(G)** Quantification of dendritic ER Ca^2+^peak release. **(H)** Corresponding quantification of fluorescence recovery 15 s after glutamate uncaging.

To examine whether ER-HaloCaMP_585_ could accurately detect physiological ER Ca^2+^release (decrease in signal) and not only uptake (increase in signal), we assessed its responsiveness in neuronal dendrites, another subcellular localization containing limited ER volume. During excitatory neurotransmission, dendrites receive extracellular glutamate inputs from other neurons, which drive Ca^2+^ entry from the extracellular space and cause Ca^2+^ release from dendritic ER^24^, ^58, 59^

To study this process, we locally released glutamate near single dendritic spines of neurons in culture using glutamate uncaging^60,61^ and examined ER Ca^2+^ responses using ER-HaloCaMP_585_. Our experiments revealed that ER-HaloCaMP_585_ responds to single-glutamate uncaging pulses with distinct ER Ca^2+^ release events and a high signal-to-noise ratio. The relative magnitude of these events was indistinguishable from those obtained with ER-GCaMP6-210 (Fig. 2E-G, see Methods).

However, when comparing the photostability of prolonged measurements of ER Ca^2+^ release, we found that signals of ER-HaloCaMP_585_ robustly recovered to the initial baseline, while ER-GCaMP6-210 showed a loss of fluorescence at the end of the experiment, suggesting reduced photostability (Fig. 2H). We also observed that ER-GCaMP6-210 exhibited a rapid decrease in fluorescence immediately after 488 nm light exposure both in dendrites and axons, which then quickly settled into a new steady-state baseline (Fig. S2A, B). While this phenomenon has been previously observed on ER-GCaMPs^58,^ its origin remains unclear. It may reflect a rapid photophysical transition, such as reversible photoswitching or photoisomerization, in which an initial exposure to blue light drives a fraction of the GFP-based molecules into a less fluorescent state, quickly reaching equilibrium^62^. Contrary to this behavior, ER-HaloCaMP_585_ maintained a stable and bright baseline signal throughout the imaging period in both axons and dendrites, exhibiting superior photostability compared to ER-GCaMP6-210 (Fig. S2A, B).

### ER-HaloCaMP fluorescence is tunable and enables far-red ER Ca^2+^ measurements

We next evaluated the usability of ER-HaloCaMP conjugated with JF_635_-HTL (ER-HaloCaMP_635_) for measuring ER Ca^2+^ levels in the far-red spectrum, which, to our knowledge, has not yet been possible. Conjugation with JF_63_s-HTL instead of JF_58_s-HTL led to excitation and emission spectra with a peak excitation at 643 nm and emission at 654 nm, confirming its usability as a far-red sensor (Supp. Fig. S3A). Using purified protein, LA-HaloCaMP_635_ presented a shift in the Ca^2+^ affinity compared to LA-HaloCaMP_585_, increasing it by approximately 2.5-fold (Table 1). ER-HaloCaMP_635_ retained a molecular brightness of 48 mM^-1^cm^-1^ which remains substantially higher than that of conventional red GECis (∼7-fold increase, Table 1). When imaged in HeLa cells, ER-HaloCaMP_635_ generated a far-red fluorescence pattern consistent with ER labeling (Fig. 3A). Upon thapsigargin addition, fluorescence decreased by about 80%, confirming its usability as a far-red ER Ca^2+^ sensor (Fig. 3B). Given the non-existence of other far-red ER Ca^2+^ sensors, we evaluated ER-HaloCaMP_635_ by comparing its AF/F changes to those of ER-GCaMP6-210 and ER-HaloCaMP_585_. The results showed that the changes detected by ER-HaloCaMP_635_ were approximately in the same range as those observed with ER-GCaMP6-210 or ER-HaloCaMP_585_ (Fig. 3C). This result supports the versatility ofER-HaloCaMP in capturing live ER Ca^2+^ dynamics in different fluorescent colors.

**Figure 3.**
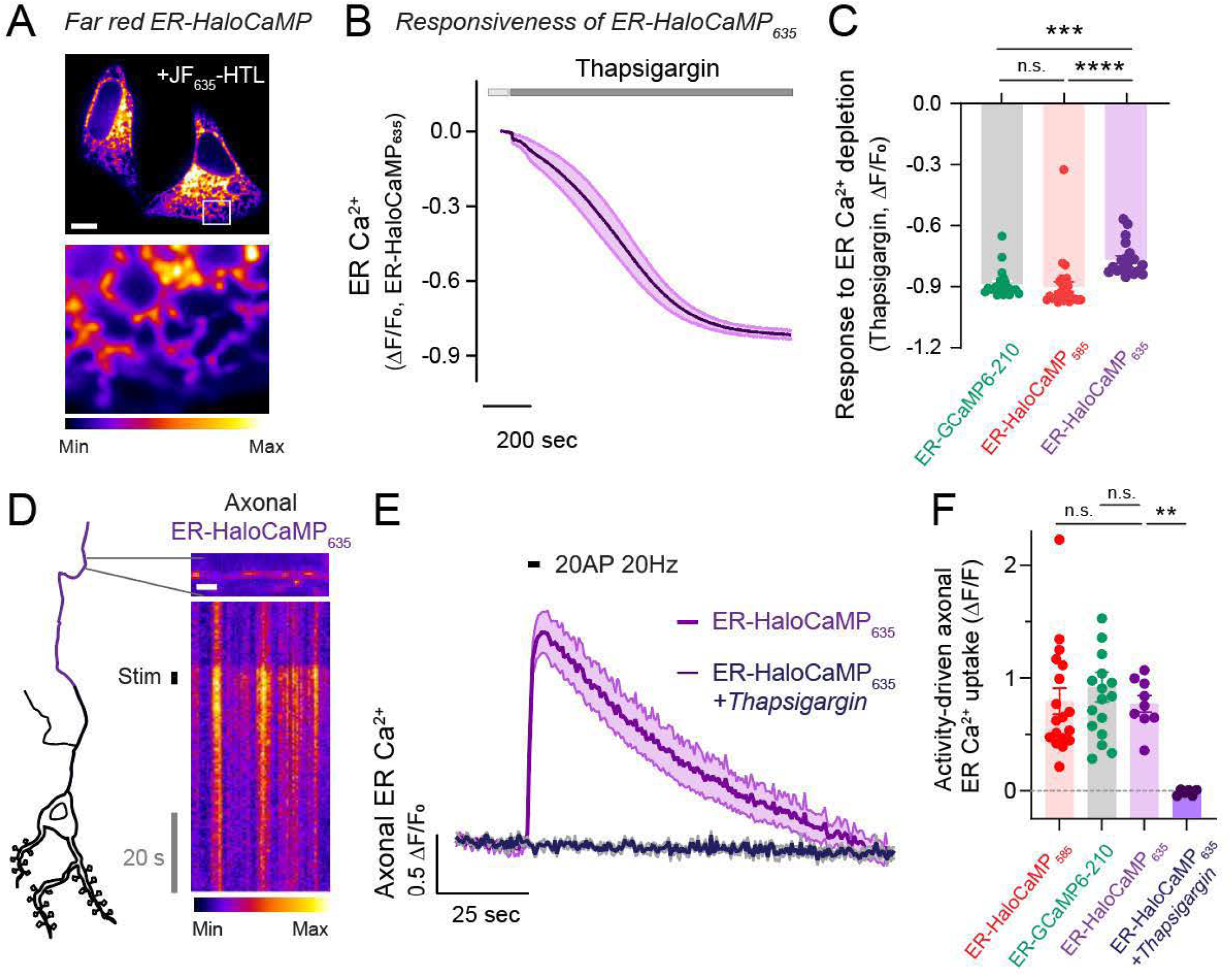
Far-red ER Ca^2+^ measurements using ER-HaloCaMP_63_5. (A) High resolution image of a HeLa. cell expressing ER-HaloCaMPla reconstituted with JF_635_-HTL shows the ER structme (pseudocolor scale below showing low to high intensity). Scale bar, 12 µm. **(B)** Average fluorescence intensity over time, upon thapsigargin treatment (I µM) in HeLa cells, measured with ER-HaloCaMP_635_. Light gray indicates cells in ‘fyrode’s solution, dark grey indicates when thapsigargin is adeed. (C) Quantification of the thapsigargin response (I µM) in HeLa. cells, measured with different sensors. **(D-F)** Axonal ER Ca^2+^ responses to 20 AP (20 Hz) stimulation, using different indicators. **(D)** On the left: schematic representation of ER-HaloCaMP_635_ expressed within a single ER tubule of axon. On the right representative image of a neuron expressing ER-HaloCaMP_635_, zoomed on an axon. Scale bar, 4 µm. Corresponding kymograph (pseudocolor scale below showing low to high intensity). (E) Corresponding fluorescence intensity over time, before and after thapsigargin treatment (I µM). (F) Quantification of activity-driven ER Ca^2+^ uptake peak response upon 20 AP (20 Hz) stimulation. Data are represented as mean ± SEM. See Supplementary Table STI for details on statistical tests and sample sizes.

We then investigated whether ER-HaloCaMP_635_ could be used to visualize and monitor ER Ca^2+^ dynamics fluorescence within the narrow ER tubules of axons. We first confirmed that the high brightness of ER-HaloCaMP_635_ enabled successful detection of axonal ER at rest (Fig. 3D). Next, we electrically stimulated neurons and observed robust axonal ER Ca^2+^ uptake. As control, signals were completely silenced by thapsigargin, confirming that the increases in fluorescence depended on ER SERCA pumps (Fig. 3E). Benchmarking against ER-GCaMP6-210 and ER-HaloCaMP_585_ showed that ER-HaloCaMP_635_ exhibited lower responses, which could be a consequence of its higher affinity (Fig. 3F). Despite this caveat, ER Ca^2+^ uptake was consistently detected in response to electrical stimulation suggesting that resting ER Ca^2+^ concentration is low enough in axons for this indicator to be useful for detecting changes in subcellular regions with low ER abundance.

### Developing a responsive and tunable red/far-red mitochondrial Ca^2+^ sensor based on BaloCaMP

Building on the robust performance of ER-HaloCaMP, we extended our strategy to develop mitochondrial Ca^2+^ sensors based on HaloCaMPs. Mitochondrial Ca^2+^ entry occurs dynamically in cells and is essential for cellular physiology, as it adjusts energy production, modulates metabolic pathways and influences cell survival and apoptosis^18^. Resting free Ca^2+^ levels in mitochondria are higher than in the cytosol, yet remain in the hundreds of nanomolar rather than micromolar range^11,63-65^. Previous studies have shown that low-affinity mitochondrial Ca^2+^ sensors exhibit inadequate responses to capture physiological stimuli in HeLa cells^19^ and neurons^66^. In contrast, adapted sensors originally developed for cytosolic Ca^2+^ measurements give robust mitochondrial responses^11, 66^. We targeted HaloCaMPla^40^ to mitochondria using tandem mitochondrial targeting sequences from cyclooxygenase 8^11^ and fused it with mTagBFP2 to create Mito-HaloCaMP (Fig. 4A). We expressed Mito-HaloCaMP in HeLa cells and conjugated it with JF_585_-HTL, which generated a bright mitochondrial pattern in HeLa cells that upon histamine addition reported a robust change in mitochondrial Ca^2+^ (Fig. 4B). To assess the efficacy ofMito-HaloCaMP_585_, we compared it against Mito^4^x-jRCaMPlb^11,67^ by measuring their respective AF/F_0_ responses and brightness under identical conditions. Our results demonstrated that Mito-HaloCaMP_585_ not only exhibited significantly larger AF/F changes but was also significantly brighter, indicating enhanced sensitivity and improved signal-to-noise for monitoring mitochondrial Ca^2+^ dynamics in HeLa cells (Fig. 4B-D).

**Figure 4.**
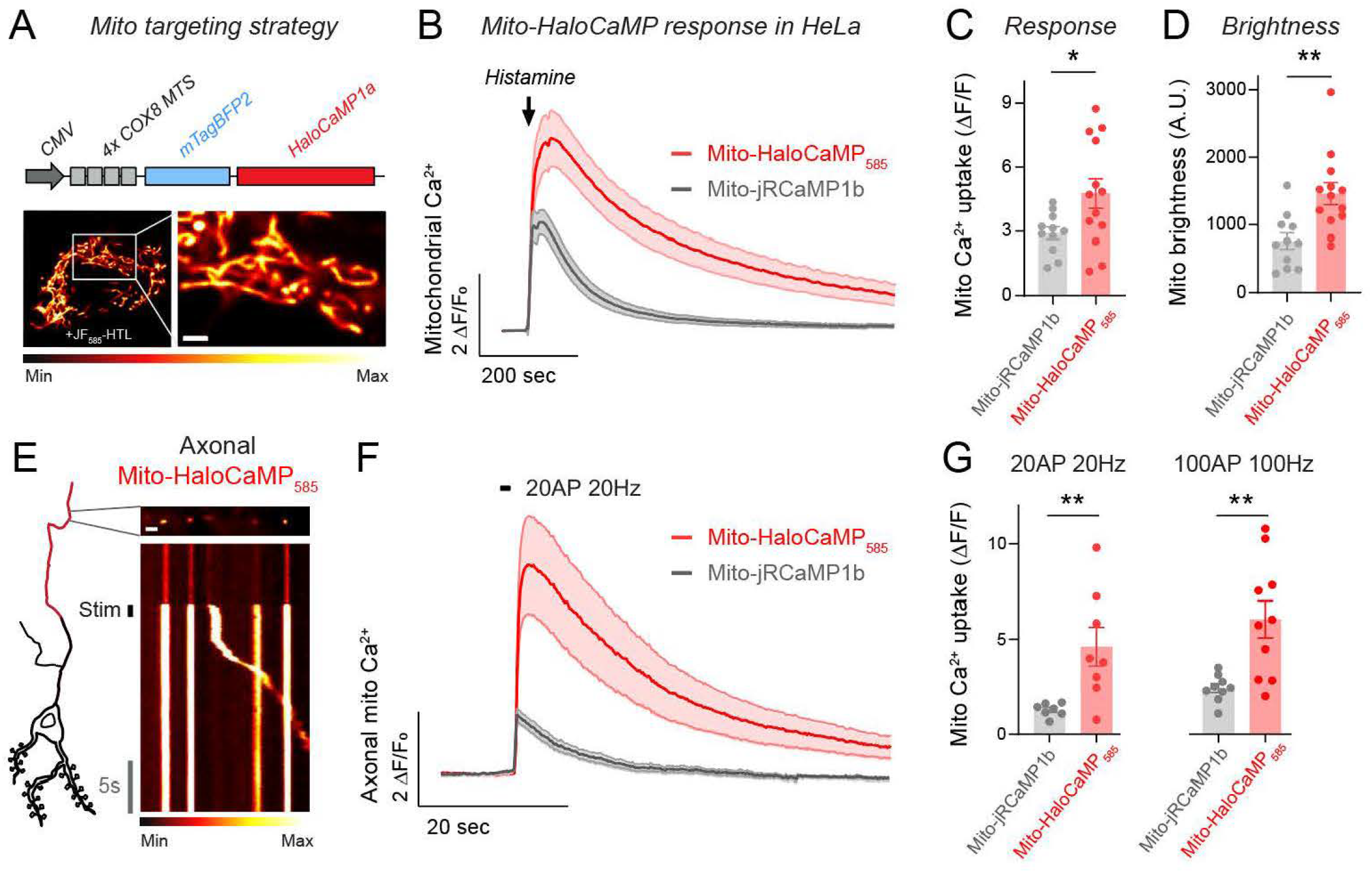
Mito-HaloCaMP is a tunable and highly responsive mitochondrial Ca^2+^ sensor. Top: targeting scheme for expression in the mitochondria by adding 4 times COX8 mitochondrial targeting sequence (MTS). Bottom: high resolution image of a HeLa cell expressing Mito-HaloCaMP reconstituted with JF_585_-HTL shows the mitochondrial structures (pseudocolor scale below showing low to high intensity). Scale bar, 2 µm. **(B)** Average fluorescence intensity over time, upon histamine treatment (10 µM), measured with red mitochondrial calcium indicators in HeLa cells. (C) Corresponding quantification of the peak response. **(D)** Comparison of relative brightness of mito-RCaMPlb and mito-HaloCaMP_585_ using identical illumination and detection conditions in intact HeLa cells. (E) On the left: schematic representation of mito-HaloCaMP_585_ expressed within a single axon. On the right representative image of a neuron expressing mito-HaloCaMP_585_, zoomed on an axon. Scale bar, 4 µm. Corresponding kymograph (pseudocolor scale below showing low to high intensity). **(F)** Corresponding average fluorescence intensity over time upon 20 AP (20 Hz) stimulation, measured with red mitochondrial calcium indicators in axons of cortical neurons. (G) Quantification of activity-driven mitochondrial Ca^2+^uptake peak response upon 20 AP (20 Hz) stimulation or 100 AP (100 Hz) with red mitochondrial calcium indicators.

In non-excitable cells and neuronal somata, mitochondria are easily detectable and form networks of elongated structures. However, in axons their morphology is significantly reduced and they appear as small rounded structures averaging approximately 1 µm in length, thereby making their detection significantly more challenging. We next used Mito-HaloCaMP_585_ to identify axonal mitochondria and electrically stimulated neurons to fire 20AP at 20 Hz as before (Fig. 4E). During neuronal firing, we observed that Mito-HaloCaMP_585_ exhibited robust mitochondrial Ca^2+^ responses, generating fluorescence changes that were 3.5-fold greater than those obtained with Mito-jRCaMPlb under identical conditions (Fig. 4F). Similarly, when stimulating neurons with lO0AP fired at 100 Hz, Mito-HaloCaMPsss still reported significantly larger changes (Fig. 4F,G). These findings indicate that Mito-HaloCaMP_585_ represents a significant advancement in mitochondrial Ca^2+^ imaging technology, providing a brighter and more responsive tool for red mitochondrial Ca^2+^ measurements.

We next evaluated the usability of Mito-HaloCaMP conjugated with JF_635_-HTL (Mito-HaloCaMP_635_) for measuring mitochondrial Ca^2+^ dynamics in the far-red spectrum, which, to our knowledge, has not yet been possible. We reasoned that the ability of measuring organellar Ca^2+^ handling in the far-red spectrum should enable quantifying Ca^2+^ dynamics in the different subcellular compartments involved in ER-mitochondria Ca^2+^ transfer, enabling the simultaneous detection of Ca^2+^ in three independent subcellular locales. We used histamine as a stimulus to drive ER Ca^2+^ release and simultaneously quantified Ca^2+^ dynamics in ER, cytosol and mitochondria of HeLa cells (Fig. SA). These experiments were performed in the absence of extracellular Ca^2+^ to ensure that all observed signals originate from the ER, thereby allowing precise quantification of how much ER Ca^2+^ is released into the cytosol versus taken up by mitochondria (see Methods). We examined the timing of Ca^2+^ changes in each compartment using the cytosolic peak as a reference point and we found that ER Ca^2+^ release occurred earlier, while mitochondrial Ca^2+^ signals lagged several seconds behind (Fig. 5B, C). These results confirm that Ca^2+^ release from the ER is the initiating event that drives subsequent cytosolic and mitochondrial Ca^2+^ accumulation. Single-cell measurements revealed that the amplitude of both cytosolic and ER Ca^2+^ correlated with the amplitude of mitochondrial Ca^2+^ increases (Fig. 5D; Fig. S3B, C). However, we observed significant variability, finding large differences in ER-mitochondrial Ca^2+^ transfer in between different cells (Fig, SE, top panel; Fig. S3A, B). As a control, we confirmed that differences did not arise from saturation of the sensors during histamine responses, as we used ionomycin at the end of every experiment to obtain the maximal response of each sensor (Fig. S3D-G). To quantify ER-mitochondrial Ca^2+^ coupling, we leveraged our simultaneous quantification of ER, mitochondria and cytosol to quantify in individual cells the distribution of ER-derived Ca^2+^ into mitochondrial and cytosolic compartments (Fig. SE, bottom; see Methods). Our results show that as ER Ca^2+^ release increases, cytosolic Ca^2+^ increases and the coupling to mitochondrial Ca^2+^ entry increases cooperatively. This indicates that ER-mitochondria Ca^2+^ transfer is not a linear process but it is dynamically adjusted with variable ER Ca^2+^ release events. This novel strategy to monitor ER-mitochondria Ca^2+^ transfer provides a robust technical framework to explore the fitness of this process in different cell types and cell states in health and disease.

**Figure 5.**
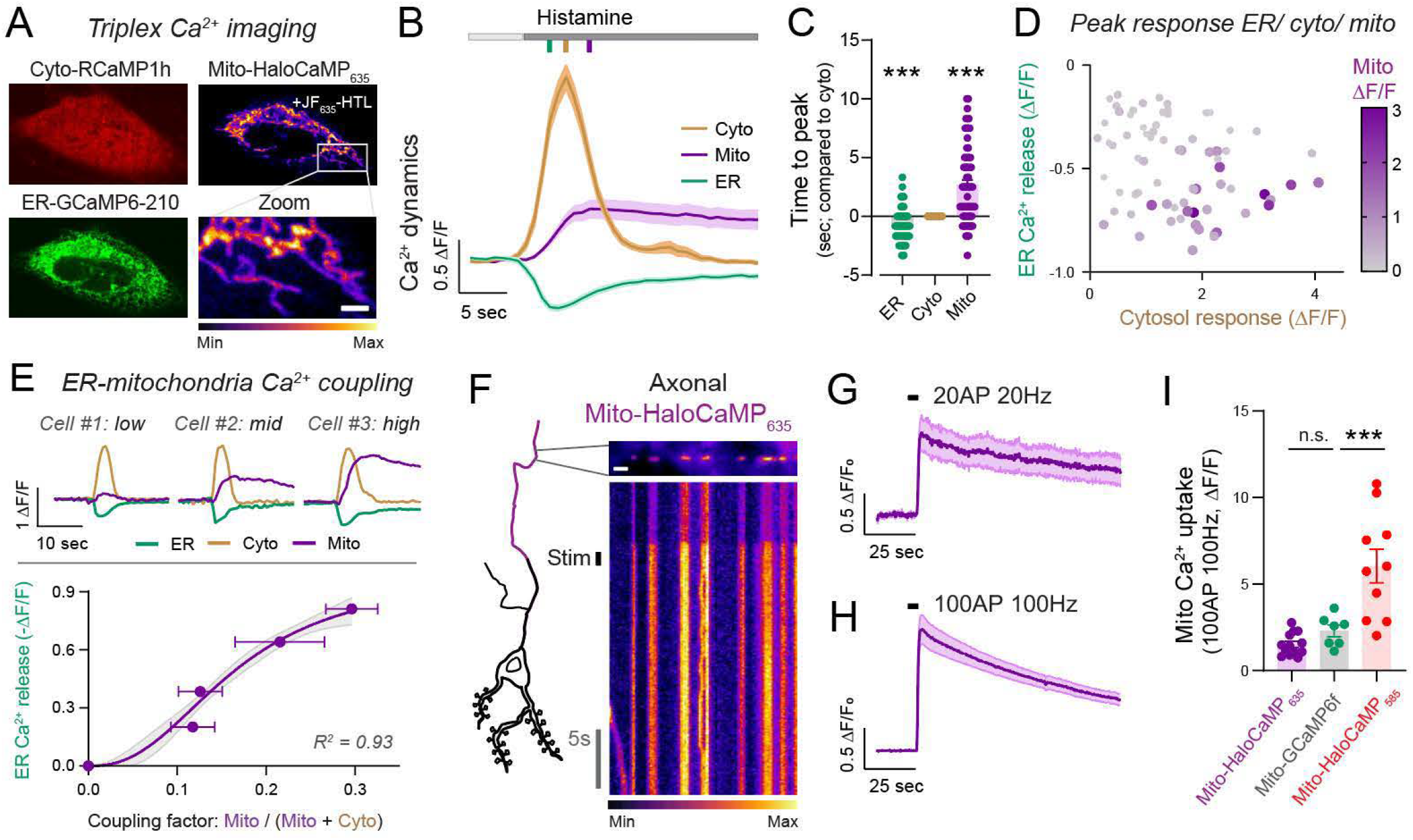
Far-red mit(rHaloCaMP imaging enables multiplexing Ca^2+^ signaling. **(A)** HeLa cell expressing cytosolic RCaMPlh (red), ER-GCaMP6-210 (green) and MitoHaloCAMP_635_. Zoomed image shows individual mitochondria. Intensity bar: low to high fluorescence. Scale bar, 2 µm. **(B)** Averaged responses to histamine (dark grey) in the cytosol (cyto, brown), mitochondria (mito, pwple), and endoplasmic reticulum (ER, green). Vertical colored bars indicate the time at which each organelle response is maximal (C) Time-to-peak Ca^2+^ responses in each compartment, calculated relative to the cytosolic peak set to t=O. **(D)** Correlation between ER, cytosolic and mitochondrial Ca^2+^ signals during histamine. Each dot represents a single cell. Dot size and color intensity are scaled to reflect mitochondrial ΔF/F. **(E)** Top panel: single cell examples showing low, medium and high mitochondrial Ca^2+^ responses for similar ER and cytosol Ca^2+^ dynamics. Lower panel: quantification of ER-mitochondrial Ca^2+^ transfer during histamine. Mitochondria and cytosol peak responses were used to calculate the coupling factor (see Methods). The line represents a fit to a Hill curve, and the gray shading indicates the 99% confidence interval of the fit. R2(coefficient of determination) = 0.93. **(F-1)** Axonal Mito-HaloCaMP_635_ Ca^2+^ responses to 20 AP (20 Hz) and 100 AP (100 Hz) stimulation (F) On the left: schematic representation of Mito-HaloCaMP_635_ expressed within a single axon. On the right representative image of a neuron expressing mito-HaloCaMP_635_, zoomed on an axon. Scale bar, 4 µm. Corresponding kymograph of response, scale bar indicates 5 sec. Pseudocolor scale shows low to high intensity. **(G)** Average fluorescence intensity over time upon 20 AP (20 Hz) stimulation. **(H)** Average fluorescence intensity over time upon 100 AP (100 Hz) stimulation. **(I)** Quantification of activity-driven mitochondrial Ca^2+^ uptake peak response upon 100 AP (100 Hz) measured with Mito-HaloCaMP635, Mito-GCaMP6f and Mito-HaloCaMP585. Data are represented as mean ± SEM. See Supplementary Table STl for details on statistical tests and sample sizes.

Lastly, we examined the capabilities of Mito-HaloCaMP_635_ to detect mitochondrial Ca^2+^ uptake in single mitochondria of the axon during neuronal activity (Fig. SF). We electrically stimulated neurons to fire 20AP at 20 Hz or lOOAP at 100 Hz as before and observed that Mito-HaloCaMP_635_ exhibited robust mitochondrial Ca^2+^ responses (Fig. 5G, H). As no far-red mitochondrial Ca^2+^ sensors currently exist, we compared Mito-HaloCaMP _635_ to Mito-GCaMP6f, the gold standard in the field. We found that Mito-GCaMP6f responses were indistinguishable from those of Mito-HaloCaMP_635_, demonstrating its high sensitivity (Fig. 5I). We also compared these data to Mito-HaloCaMP _585_, which performed significantly better than Mito-GCaMP6f (Fig. 5I). These results show that Mito-HaloCaMP enables robust detection of mitochondrial Ca^2+^ changes in red and far-red even in challenging imaging conditions, such as detecting fluorescence changes in single small axonal mitochondria.

### ER-HaloCaMP Ca^2+^ measurements in brain tissue of different species

Lastly, to extend the applicability of organellar HaloCaMPs beyond cultured cells, we next assessed the ability of ER-HaloCaMP_585_ to report ER Ca^2+^ dynamics in *ex vivo* brain preparations from rats and Drosophila. We first expressed ER-HaloCaMP in individual CA3 pyramidal neurons in organotypic rat hippocampal slices (Fig. 6A). After labeling the sensor with JF_585_-HTL, we triggered and quantified somatic ER Ca^2+^ release via DHPG (3,5-dihydroxyphenylglycine), a group I metabotropic glutamate receptor agonist that activates the production of inositol 1,4,5-trisphosphate (IP_3_) and the release of Ca^2+^

**Figure 6.**
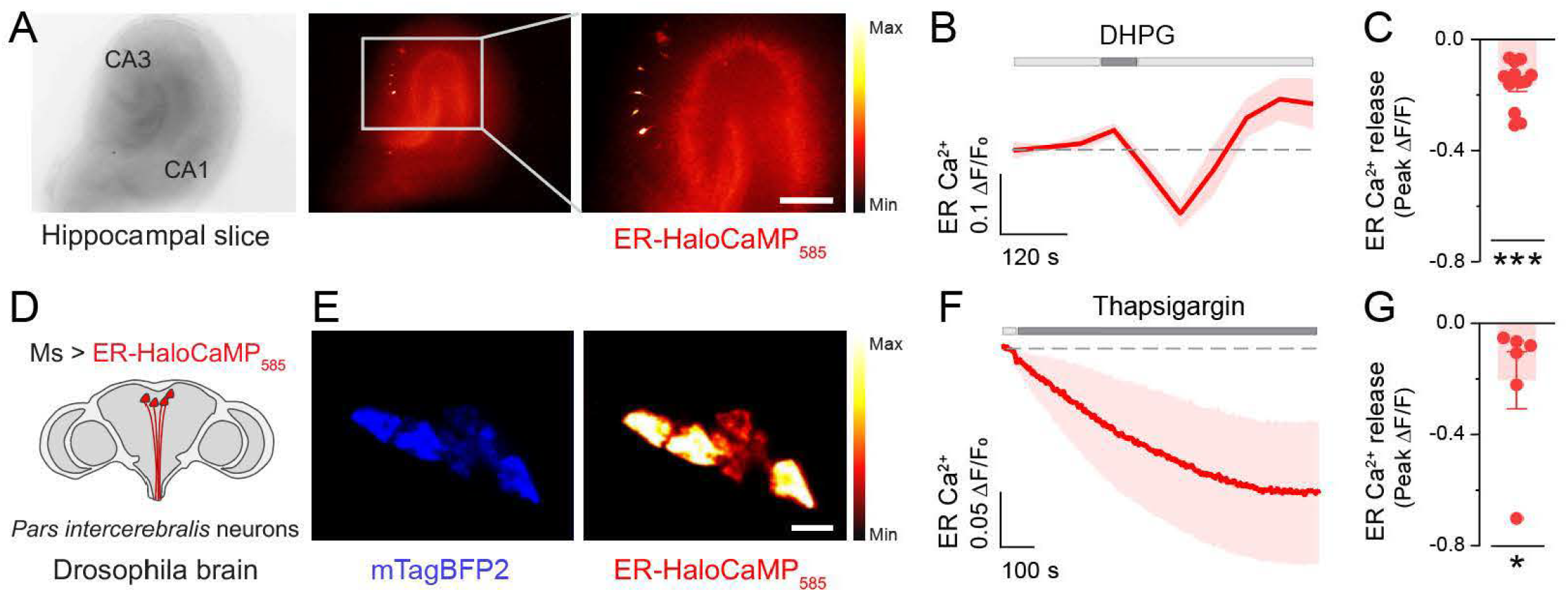
Visualization of ER Ca^2+^ dynamics in rat and fly brain tissue using ER-BaloCaMP. **(A)** Representative images of an organotypic hippocampal slice with individual CA3 neurons expressing ER-HaloCaMP_585_. A magnified view of neurons is shown on the right. Scale bar, 300 µm. **(B)** ER Ca^2+^ decreases after a pulse of 1 min application of 50 µM DHPG and overshoots later during washout. Line and shading are mean ± SEM. (C) Quantification of ER Ca^2+^ release by DHPG in individual neurons using ER-HaloCaMP_585_. **(D)** Scheme representing the fly brain and the pars intercerebralis region in which Myosupressin (Ms) neurons are located. ER-HaloCaMP is expressed in Ms neurons, denoted by Ms > ER-HaloCaMP_585_. (E) Representative 2-photon images of mTagBFP2 and ER-HaloCaMP reconstituted with JF_585_-HTL. (F) ER Ca^2+^ depletion by thapsigargin in Ms neurons using ER-HaloCaMP_585_. (G) Quantification of ER Ca^2+^ depletion by thapsigargin in individual fly brains using ER-HaloCaMP_585_. See Supplementary Table STl for details on statistical tests and sample sizes.

from internal ER stores^68,69^. Brief application of DHPG led to a transient reduction in neuronal ER Ca^2+^ levels, showing that ER-HaloCaMP_585_ can successfully report ER Ca^2+^ dynamics in thick samples using two-photon microscopy (Fig. 6B). Next, we evaluated ER-HaloCaMP_585_ functionality in Drosophila neurons. Specifically, we targeted sensor expression to Myosuppressin (Ms) neurons within the *pars intercerebralis*, an evolutionarily conserved neuroendocrine center of the fly brain (Fig. 6C). Following labeling with JF_58_,-HTL, ER-HaloCaMP fluorescence was robustly detectable within the somata of Ms neurons (Fig. 6D). Treatment with thapsigargin resulted in a clear and progressive decrease in ER Ca^2+^ levels in Ms neurons as reported by the decrease of ER-HaloCaMP_585_ fluorescence (Fig. 6E), while the fluorescence of mTagBFP2 remained unaltered (Fig. S4A-C). These results demonstrate that ER-HaloCaMP can report ER Ca^2+^ dynamics across tissues from evolutionarily distant species, validating its versatility as a tool for studying ER Ca^2+^ signaling in intact tissue.

## Discussion

The interplay between cytosolic and organellar Ca^2+^ pools orchestrates a myriad of cellular responses essential for cellular function and survival2.^70^ and alterations in organellar Ca^2+^ signaling are heavily associated with disease^18, 71^, ^75^ However, much of our understanding of organellar Ca^2+^ fluxes has relied on indirect approaches or on monitoring Ca^2+^ in different compartments in separate experiments due to limited multiplexing capabilities^24,76-78^. For example, when studying ER-mitochondria Ca^2+^ transfer, dual-compartment Ca^2+^ imaging comparing cytosol and mitochondria, or cytosol and the ER, have been used to infer the relationship between cytosolic, ER and mitochondrial Ca^2+^; ^11, 24, 77^ While these are highly valuable approaches, our triplex ER-cytosol-mitochondria Ca^2+^ measurements showed there is large cell-to-cell variability, suggesting that quantifying ER-mitochondria Ca^2+^ transfer in different populations of cells using only 2 sensors may complicate interpretations^15, 18^, ^79^, ^80^ The ability to image ER and mitochondrial Ca^2+^ using the far-red part of the spectrum unlocks distinguishing the relationship between the three compartments in single cells, enabling a precise quantification of ER-mitochondria Ca^2+^ transfer that takes into account the amplitude of cytosolic signals. Triplex imaging also enables the dissection of the temporal sequence of Ca^2+^ fluxes, revealing that during histamine stimulation, Ca^2+^ is first released from the ER, subsequently peaks in the cytosol, and is then taken up by mitochondria. Our experiments also reveal that ER Ca^2+^ release favors ER-mitochondria Ca^2+^ coupling in a cooperative manner. These findings in intact cells support the current model established in isolated mitochondria and permeabilized cells, in which opening of the Mitochondrial Ca^2+^ Uniporter (MCU) is controlled by cooperative Ca^2+^ binding to accessory Mitochondrial Calcium Uptake (MICU) proteins^81 82^ Overall, these experiments reveal the importance of simultaneous single-cell measurements of different compartments when studying organellar Ca^2+^ signaling and reveal the greatpotential of far-red imaging using organellar HaloCaMPs to explore complicated biological questions.

In addition to expanded multiplexing capacities, the development of ER-HaloCAMP and Mito-HaloCaMP presents 2 important advancements. First, they show significantly improved brightness and responsiveness in both ER and mitochondrial Ca^2+^ measurements in the red spectrum: the improved brightness of our organellar sensors likely reflects the higher extinction coefficients and quantum yields of synthetic dyes^40, 41^., as was shown when cytosolic HaloCaMP1 _35_ was compared to other protein-based fluorescent Ca^2+^ sensors, such as GCaMP6s^40^. Enhanced brightness is of particular importance when measuring Ca^2+^ in organelles because quantifying Ca^2+^ dynamics in individual mitochondria^11,13,14^ or in narrow ER tubules^12^ is a demanding task that only a few sensors are able to achieve. Second, they present improved photostability compared to the current best-in-class sensors: we found that upon blue light illumination, ER-GCaMPs underwent a rapid decrease in fluorescence that reaches stability in a few seconds, as shown previously5^8^. Such photoinactivation behavior has already been reported in some cytosolic GCaMPs, including preliminary versions of the original GCaMP, as well as GCaMP7c and jGCaMP8 variants^8^3-8s. Recent work on jGCaMP8 proposed that such behavior is the consequence of reversible photoswitching, likely caused by cis-trans isomerization of the GFP chromophore, which shifts the sensor to a non-fluorescent state independent ofCa^2+^ binding^83^. This indicates that ER-GCaMPs likely suffer from the same problem, which distorts signals independently of ER Ca^2+^ changes and confounds the quantification of rapid Ca^2+^ transients at the start of the imaging period. Because ER-HaloCaMP uses an external synthetic fluorophore rather than a circularly permuted GFP, it cannot undergo the cis-trans isomerization that promotes photoswitching. Thus, ER-HaloCaMP shows significantly more stable fluorescence signals than ER-GCaMP while retaining similar Ca^2+^ responsiveness, making it the ideal choice for endoplasmic reticulum Ca^2+^ imaging.

Organellar HaloCaMPs also demonstrate robustness across diverse biological models (cultured cell lines, primary neurons and intact tissues from different species) and are compatible with both one- and two-photon excitation This broad compatibility across model and imaging systems underscores the versatility of the HaloCaMP toolkit for investigating organellar Ca^2+^ signaling in diverse biological contexts and experimental conditions, which will facilitate their use in future research. Overall, we show that ER-HaloCaMP and Mito-HaloCaMP present improved properties that make them the preferred choice in several Ca^2+^ signaling paradigms in cell lines and primary cells. Thus, our expanded palette of organellar Ca^2+^ sensors unlocks new possibilities for the study of organellar Ca^2+^ signaling across fields, which will facilitate our understanding of organellar cell physiology in health and disease.

## Material and methods

## Lead contact

Further information and requests for resources and reagents should be directed to and will be fulfilled by the lead contact, Dr. Jaime de Juan-Sanz.

## Resources Table

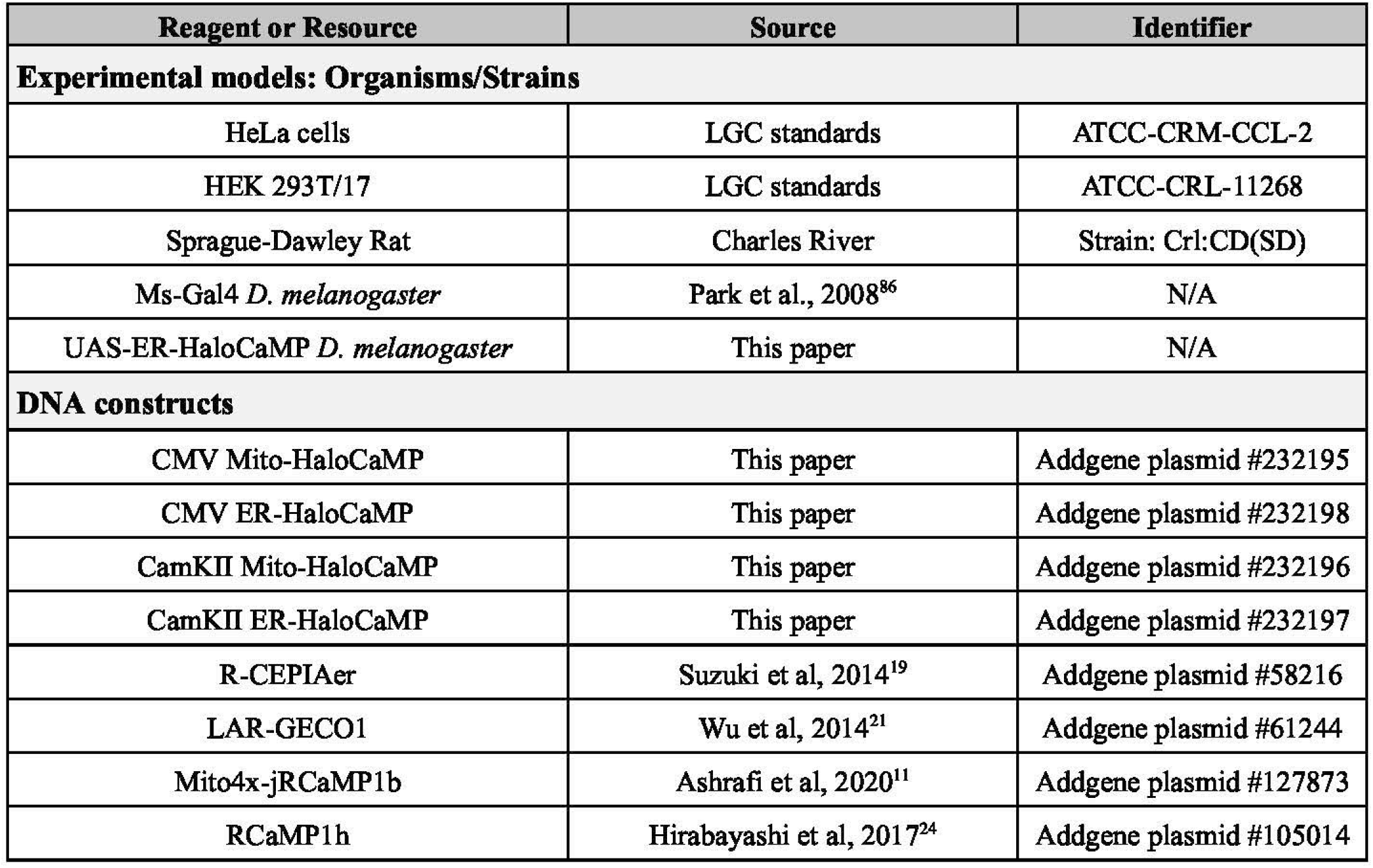

Molecular biology for generating the low affinity HaloCaMPs

Synthetic DNA oligonucleotides were purchased from Integrated DNA technologies. *QS* high fidelity DNA polymerase (New England Biolabs) wu used for all PCR amplifications. Isothermal assembly reactions were performed with a NEBuilder HiFi kit (New England Biolabs). Small scale DNA isolation were performed with QIAprep Spin Miniprep Kit (Qiagen). The pRSET vector backbone was acquired from Life Technologies. Inserts and vector backbones were amplified by PCR amplification. Vector backbones and inserts were assembled by isothennal assembly with 10-30 base pair overlap, and sequence verified by Sanger sequencing (Azenta Life Sciences) or by nanopore filll-plasmid sequencing (Plasmidsaurus).

### Protein npnssfon and parifleation

For expression and purification of proteins, T7 express (New England Biolabs) were tnmsformed with pRSET plasmids encoding for the protein of interest The bacteria were grown in auto-induction media using the Studier methodn^87^ with antibiotics at 30 °C for 48 h shaking at 200 rotations per miute (r.p.m.). Cell pellets were collected by centrifugation, lysed in TRIS Buffered Saline (TBS; 19.98 mM Tris, 136 mM NaCl, pH 8.0), with n-Octyl-β -D-thioglucopyranoside (S g L^-1^). Aggregations were disrupted by sonication and the lysate cleared by centrifugation. Protein purification was performed on a N-tenninal poly-histidine (His_6_,) tag using HisPur Ni-NTA resin (Thennofisher Scientific), acoording to manufacturer’s recornmendatio Purified proteins were buffer exchanged into TBS using Amicon concentration filters (Merck). Protein aliquots were stored at 4 °C.

### Calefam titrations in purified protein

To determine the calcium affinity and cooperativity of the low affinity HaloCaMP sensom, calcium titrations were performed in a buffer system made from CaNTA and NTA prepared using the pH titration method descried by Tsien and Pozza^88^ mixed in specific ratios to generate known free calcium concentrations. The free calcium ooncentration was calculated assuming the dissociation oonstant of NTA for Ca^2+^ 67 μ M. HaloCaMP proteins were pre-labeled with a limiting dye-HaloTag ligand (20 pM protein, 10 μM JF_585_-HTL). 2 μL of the pre-labeled protein-dye oonjugate was diluted into 98 μL of pre-mixed solutions of Ca-NTA in black 96-well plates. Fluorescence intensities were read on a plate reader (Tecan Spark 20M). Fluorescence intensity was measured at 26 °C and 37 °C. For HaloCaMPs with JF_585_-HTL excitation was 590 nm and emission 620 nm. For GCaMPs excitation was 484 nm and emission wu *S*10 nm. All bandwidths were set to 5 nm. Changes in fluorescence in addition of Ca^2+^ were calculated in Microsoft Excel. The fluorescence (y) was plotted against the free calcium ooncentration (x) and a four-parameter dose-response curve (variable slope) using GraphPad Prism vl0 software was fit where (a) is the value of fluorescence at the bottomof the curve, (b) is the value of fluorescence at the top of the cuve, (EC_50_) is the ooncentration of agonist that givesa response halfwaybetweenbottom and the top, and (h) is the hillor cooperative coefficient.

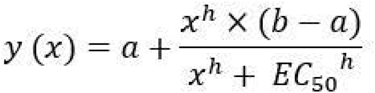

### Seqaenee of LA-HaloCaMPl

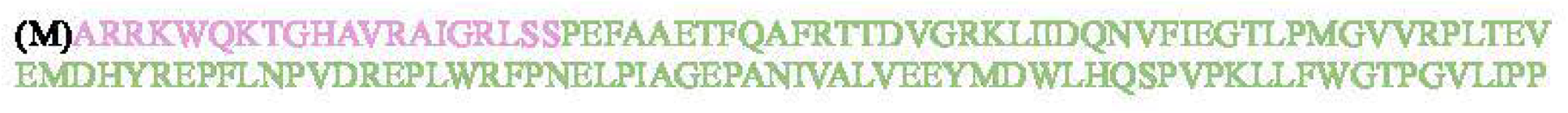
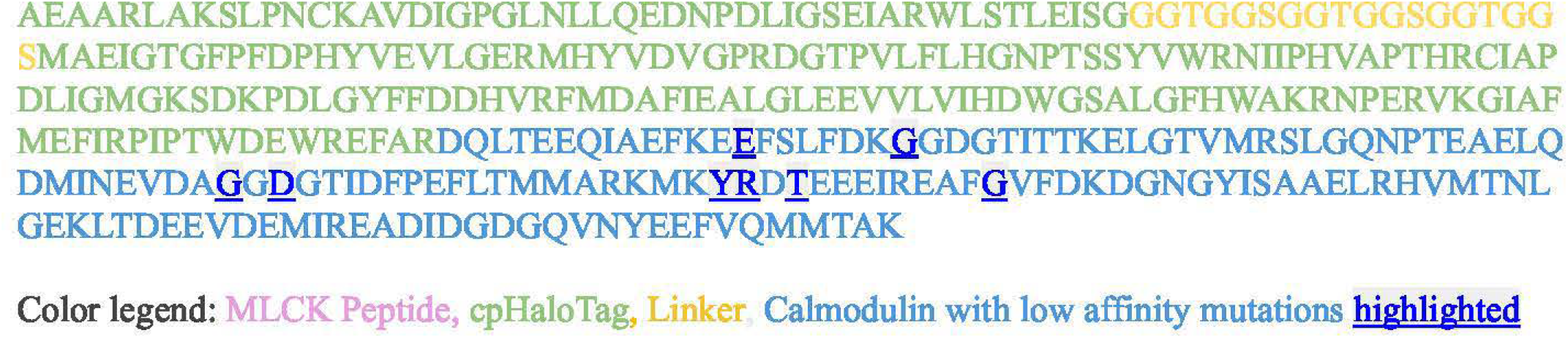

### Animals

The rats of either sex used in this study to prepare primary cultures were of the Sprague-Dawley strain, Crl (SD), bred by Charles River Laboratories following the international genetic standard protocol (IGS). All experiments conducted at the Paris Brain Institute strictly followed the guidelines set by the European Directive 2010/63/EU and the French Decree n° 2013-118 for the protection of animals used in scientific research. In the case of organotypic slices, rats were housed and bred at the University Medical Center Hamburg-Eppendorf (UKE). All procedures complied with German animal welfare regulations (Tierschutzgesetz der Bundesrepublik: Deutschland, TierSchG) and conformed to Directive 2010/63/EU. Experimental protocols were approved by the Behorde fiir Justiz und Verbraucherschutz (BN), Lebensmittelsicherheit und Veterinärwesen, Hamburg.

Flies were raised on a standard cornmeal/agar diet (6.65% cornmeal, 7.15% dextrose, 5% yeast, 0.66% agar, supplemented with 2.2% nipagin and 3.4mL/L propionic acid). All experimental flies were kept in incubators at 70% humidity and on a 12h light/dark cycle, at 25°C. Flies were transferred to fresh vials every 2 days, and fly density was kept to a maximum of 20 flies per vial.

### Generation of UAS-ER-HaLo-CaMP::BFP transgenic line

The ER-HaloCaMP fragment was PCR amplified using the primers (5’-aagatcctctagaggtacccTTAGAGTTCATCCTTGCC-3’) and (5’-actctgaatagggaattgggATGGGACTGTTGTCTGTTC-3’). The fragment was then cloned in the open pUASTattB vector through site-directed mutagenesis using the kit Q5-site directed mutagenesis (NEB E0554S). The pUASTattB vector was opened with restriction digestion using the restriction enzymes EcoRI-HF (NEB R3101S) and Xhol (NEB R0146S). The resulting pUASTattB-ER-HaloCaMP plasmid was used to establish transgenic lines through cpC-31 integrase mediated recombination (BestGene Inc.). The landing attP site used was VK37 PBac{yellow[+]-attP-3B}VK00037, BDSC Stock 9752.

### Primary rat co-culture of postnatal cortical neurons and astrocytes

Primary co-cultures of cortical neurons and astrocytes were obtained as previously described with small modifications^89^. P0 to P2 rats of mixed gender were sacrificed, and their brains were dissected in a cold HBSS-FBS solution (lX HBSS + 20% Fetal Bovine Serum) to isolate the cerebral cortexes. These were then cut into small pieces for digestion and dissociation. The tissue was washed twice with 1X HBSS-FBS and lX HBSS before being incubated in a trypsin-based digestion solution containing DNAse I (Merck, D5025) for 15 minutes at room temperature. Trypsin (Merck, T1005) was neutralized with HBSS-FBS solution, followed by two washes with lX HBSS-FBS and two with lX HBSS. The tissue was then transferred to a dissociation solution (lX HBSS, 5.85mM MgSO_4_) and dissociated into single cells by gentle pipetting. The cells were centrifuged at 13,000 rpm for 10 minutes at 4°C, and the pellet was resuspended in lX HBSS solution. After a second centrifugation, the pellet was resuspended in a homemade warmed plating media consisting of MEM (Thermo Fisher Scientific, 51200038) supplemented with 20 mM Glucose (Merck, 08270), 0.1 mg/ml transferrin (Merck, 616420), 1% GlutaMAX (Thermo Fisher Scientific, 35050061), 24 µg/ml insulin (Merck, 16634), and 10% FBS (Thermo Fisher Scientific, 10082147) and 2% N-21 (Bio-techne, AR008). Finally, cells were plated in sterilized cloning cylinders (Merck, C7983; 38,000 cells per cylinder) attached to coverslips (Diameter= 25 mm, Warner Instruments, 640705) that had been pre-coated with 0.1 mg/ml poly-ornithine (Merck, P3655). After 3-4 days, the neuronal media was replaced with a homemade feeding media, similar in composition to the plating media but containing *5%* FBS and 2 µM cytosine P-d-arabinofuranoside (Merck, C6645) to inhibit glial growth. The primary co-cultures of cortical neurons and astrocytes were maintained at 37°C in a humidified incubator with *95%* air/5% CO2 until the imaging experiments, which were performed from days in vitro (DIV)12 to DIV21.

For cultures used in glutamate uncaging experiments shown in Figure 2E-H, conditions were as follows: hippocampal regions were dissected in ACSF containing (in mM) 124 NaCl, 5 KCl, 1.3 MgSO_4_:7H_2_O, 1.25 NaH_2_PO_4_:H_2_O, 2 CaCl_2_, 26 NaHCO_3_, and 11 Glucose (stored at 4°C) and stored in hibernate E buffer (BrainBits LLC, stored at 4 °C). Dissected hippocampi were dissociated using Papain Dissociation System (Worthington Biochemical Corporation, stored at 4 °C) with a modified manufacturer’s protocol. Briefly, hippocampi were digested in papain solution (20 units of papain per ml in 1 mM L-cysteine with 0.5 mM EDTA) supplemented with DNase I (final concentration *95* units per ml) and shook for 30-60 min at 37°C, 900 rpm. Digested tissue was triturated and set for 3 min, following which the supernatant devoid of tissue chunks was collected. The supernatant was centrifuged at 300 ref for 5 min and the pellet was resuspended in resuspension buffer (1 mg of ovomucoid inhibitor, 1 mg of albumin, and *95* units of DNase I per ml in EBSS). The cells were forced to pass through a discontinuous density gradient formed by the resuspension buffer and the Ovomucoid protease inhibitor (10 mg per ml) with bovine serum albumin (10 mg per ml) by centrifuging at 600 rpm for 6 min. The final cell pellet devoid of membrane fragments was resuspended in Neurobasal-A medium (Gibco, stored at 4 °C) supplemented with Glutamax (Gibco, stored at -20 °C) and B27 (Gibco, stored at -20 °C). Cells were plated on poly-D-lysine coated coverslips mounted on MatTek dishes at a density of 30000-50000 cells/cm^90^. Cultures were maintained at 37 °C and 5% CO2 with feeding every 3 days using the same medium until transfection. Transfections were performed 12 days after plating by magnetofection using Combimag (OZ biosciences, stored at 4 °C) and Lipofectamine 2000 (Invitrogen, stored at 4 °C) according to manufacturer’s instructions.

### Preparation of Organotypic Hippocampal Slices

Organotypic hippocampal slices were prepared from Wistar rats of both sexes at postnatal days 5-7. Following dissection, the hippocampi were sectioned into 400 µm slices using a tissue chopper and transferred onto a porous membrane (Millicell CM, Millipore) for culture. Slices were maintained at 37 °C in an atmosphere of *5%* CO2 in a culture medium consisting of 394 mL Minimal Essential Medium (Sigma M7278), 100 mL heat-inactivated donor horse serum (Sigma Hl 138), 1 mM L-glutamine (Gibco 25030-024), 0.01 mg/mL insulin (Sigma 16634), 1.45 mL 5 M NaCl (Sigma S5150), 2 mM MgSO_4_ (Fluka 63126), 1.44 mM CaC1_2_ (Fluka 21114), 0.00125% ascorbic acid (Fluka 11140), and 13 mM D-glucose (Fluka 49152). The culture medium was partially replaced (60-70%) twice per week. To express ER-HaloCaMP in CA3 neurons of the cultured slice, plasmids encoding ER-HaloCaMP-BFP were diluted to a final concentration of 10 ng/µL in a K-gluconate-based intracellular solution containing (in mM): 135 K-gluconate, 4 MgC1_2,_ 4 N -ATP, 0.4 Na-OTP, 10 N -phosphocreatine, 3 ascorbate, and 10 HEPES (pH 7.2). Single-cell electroporation was performed between DIV 6 and DIV 10. During the electroporation procedure, slice cultures were maintained in a pre-warmed, HEPES-bu:ffered solution containing (in mM): 145 NaCl, 10 HEPES, 25 D-glucose, 1 MgC1_2_, and 2 CaC1_2_ (pH 7.4, sterile filtered). Electroporation was carried out using an ELectroPORATOR (NPI), applying 12 voltage pulses (− 12 V, 0.7 ms) at 50 Hz. Slices were incubated for approx. 24h with JF_585_-HTL in culture medium 5-7 days after electroporation, then rinsed in the pre-warmed HEPES-bu:ffered solution before imaging. We found that #x223C; 40% of ER-HaloCaMP transfected neurons in the organotypic slices were not sufficiently labelled by incubation with JF_5_ss-HTL, presenting detectable BFP while no ER-HaloCaMP_585_ fluorescence. This could be the consequence of the limited bioavailability of JF*sss-*HTL^41^, although lack of labeling was not observed when reconstituting ER-HaloCaMP in Drosophila neurons.

### Cell line culture and transfection

HeLa and HEK cells were cultured either on poly-omithine coated glass coverslips or wells of a chambered coverslip with 8 wells and a glass bottom (CliniSciences, 80807-90). They were cultured in DMEM (Thermo Fisher Scientific, 10566016) supplemented with 10% fetal bovine serum and kept in an incubator with humidified air containing 5% CO_2_ at a temperature of 37°C. HeLa and HEK cells were transfected using Lipofectamine 2000 (Invitrogen) the day before imaging following the manufacturer’s recommendations.

### HaloCaMP labelling in cultured cells

JF dye-ligands were obtained from Dr. Lavis (Janelia Research Campus; JF_585_-HTL) or from Promega (HT 1050, Janelia Fluor 635 HaloTag Ligand; JF_635_-HTL). The JF dye-ligands obtained from Promega were incubated following the manufacturers instructions. JF dye-ligands obtained from Dr. Lavis were received in aliquots of 100 nmol and resuspended in 200 µL DMSO to yield a 500 µM stock. JF dye-ligands were then diluted in the cell culture media (either DMEM for cell lines or feeding media for primary neuronal cultures or organotypic slices) at a final concentration of 1µM. The cells are usually labelled after 30 minutes of incubation but for practical purposes, they were incubated overnight with the dye-ligand. Next day, the dye-ligand was washed in the morning before the experiment. To minimize non-specific labeling and reduce background fluorescence, a series of washing steps were performed by replacing the existing media with fresh, pre-warmed culture media, followed by a 10-minute incubation period. This washing step was repeated twice to ensure thorough removal of non-specific signals and to facilitate subsequent fluorescence measurements.

### Calcium imaging in cells and primary neurons

Live imaging assays of primary cortical neurons transfected with calcium phosphate method^91^ were conducted from DIV12 to DIV21. HEK or HeLa cells imaging assays were performed one day after transfection using lipofectamine 2000 (Thermo Fisher Scientific, 11668019). The experiments utilised a custom-built, laser-illuminated epifluorescence microscope (Zeiss Axio Observer 3) paired with an Andor i.Xon Life camera (model IXON-L-897), cooled to -70°C. Illumination was provided by fiber-coupled lasers at wavelengths of 488 and 561 nm (Coherent OBIS 488 nm and 561 nm), combined using the Coherent Galaxy Beam Combiner. Laser illumination was controlled by a custom Arduino-based circuit that synchronized imaging and illumination. Neuron-astrocyte co-cultures, HEK or HeLa on coverslips were placed in a closed bath imaging chamber for field stimulation (Warner Instruments, RC-21BRFS) and imaged with a 40x Zeiss oil objective Plan-Neofluar with a numerical aperture of 1.30 (WD = 0.21 mm). Neurons were stimulated using 1 ms current pulses between platinum-iridium electrodes in the stimulating chamber (Warner Instruments, PH-2), driven by a stimulus isolator (WPI, MODEL A382) controlled by the Arduino-based circuit. All the experiments were performed at 37°C. Temperature was kept constant using a Dual Channel Temperature Controller (Warner Instruments, TC-344C) that controlled the temperature of the stimulation chamber (Warner Instruments, PH-2) and simultaneously warmed the imaging solutions using an in-line solution heater (Warner Instruments, SHM-6). Imaging solutions were flown at 0.35-0.40 mVmin. Imaging was performed using a Tyrode’s solution composed of (in mM) 119 NaCl, 2.5 KCl, 2 CaC12, 2 MgC12, 20 Glucose together with 10 µM CNQX and 50 µM AP5, buffered to pH 7.4 at 37°C using 25 mM HEPES.

### Multiplexed confocal calcium imaging in cell lines and analysis of ER-mitochondria coupling

HeLa cells expressing ER-GCaMP6-210, cytosolic RCaMPlh and Mito-HaloCaMP were cultured in chambered coverslips with 8 wells and a #1.5H glass bottom (lbidi, 80827) and transfected using lipofectamine 2000 following manufacturer’s instructions (Thermo Fisher Scientific, 11668019) to be imaged the next day. Live imaging assays were conducted with an inverted Leica spinning disk equipped with a Yokogawa CSU-Xl module and a Hamamatsu Orea Flash 4.0 sCMOS camera. The objective used was 63x, NA 1.4. The effective sampling rate at each wavelength was approximately 0.83 Hz, given a 1.2s interval between successive images. The samples were illuminated using lasers at 488 nm, 561 nm, and 637 nm. The emission was collected through 525/50, 607/35, and 685/40 filters mounted in an automated rotating filter wheel. Imaging was performed using a Tyrode’s solution composed of (in mM) 119 NaCl, 2.5 KCl, 2 CaC1_2,_ 2 MgC1_2,_ 20 Glucose, buffered to pH 7.4 at 37°C using 25 mM HEPES.

Regions of interest (ROls) were drawn over individual cells, covering ER, cytosol and mitochondria. Background-substracted fluorescence time-courses (ΔAF/F_0_) were calculated and peak responses were extracted from each channel individually after histamine addition. Cells typically showed multiple peak responses but only the first peak was used for analysis for consistency. A modified Tyrode’s solution containing 4mM Ca^2+^ and 500 µM ionomycin was added at the end of the experiment to confirm histamine responses did not saturate any of the sensors. When plotting triplex correlations, the size of the dots representing single cells ranged from 0.01 to 0.90 in arbitrary size units. Responses > 0.90 are shown at maximal dot size. ER-mitochondria coupling was quantified by calculating, for each cell initial response, the ratio of mitochondrial ΔF/F_0_ to the sum of mitochondrial plus cytosolic ΔF/F_0_. Data points were grouped into bins of 0.15 ΔF/F_0_ along the ER ΔF/F_0_ axis and the corresponding averaged coupling factors were plotted against ER ΔF/F_0_ to assess cooperativity. To obtain averaged responses, cytosolic, ER and mitochondrial ΔF/F_0_ traces were time-aligned by setting the histamine-induced cytosolic peak tot= 0, then averaged across cells. Only cells in which the peak was higher than six times the standard deviation of the baseline were included in this analysis (65 out of 71).

### Measurement of dendritic ER-HaloCaMP and ER-GCaMP6-210 upon two-photon glutamate uncaging

For the two-photon glutamate uncaging experiments in Figure 2E-H, imaging was conducted 15-16 days after neuronal cell culture plating in a modified E4 imaging buffer containing: 120 mM NaCl, 3 mM KCl, 10 mM HEPES (buffered to pH 7.4), 4 mM CaC1_2,_ and 10 mM Glucose, lacking MgC1_2._ Imaging was performed using a custom-built inverted spinning disk confocal microscope (3i imaging systems; model CSU-Wl) attached to an Andor iXon Life 888. Image acquisition was controlled by SlideBook 6 software. Images were acquired with a Plan-Apochromat 63x/1.4 NA. Oil objective, M27 with DIC III prism, using a CSU-Wl Dichroic for 488/561 nm excitation with Quad emitter and individual emitters, at back aperture laser powers 2.00 mW (488 nm) and 2.65 mW (561 nm) for spine stimulation measurements. During imaging, the temperature was maintained at 37°C using an Okolab stage top incubator with temperature control.

For ER-HaloCaMP measurements, cytosolic GCaMP6s was co-transfected to confirm effective spine stimulation (data not shown). For ER-GCaMP6-210 measurement, neurons were co-transfected with PSD95-mCherry to identify spines for stimulation. Before glutamate uncaging, neurons were treated with 1 µM TTX (Citrate salt, mM stock made in water, Abeam, ab1200552) and 2 mM 4-Methoxy-7-nitroindolinyl-caged-L-glutamate (MNI caged glutamate, Tocris Bioscience 1490, 100 mM stock made in modified E4 buffer) in modified E4 buffer lacking Mg^2+^ (see above). Glutamate uncaging was performed using a multiphoton-laser 720 nm (Mai TAI HP) and a Pocket cell (ConOptics) for controlling the uncaging pulses. Uncaging protocols of 6 uncaging pulses at 0.25 Hz with 100 ms pulse duration per pixel at 7.2 mW back aperture laser power was used.

To measure ER-HaloCaMP and ER-GCaMP6-210 response, ∼2 µm regions of interest (ROI) were drawn at the base of the stimulated spine. The average intensities were background subtracted using the intensity measured from an adjacent background area. For each successive time point during and after stimulation, the normalized intensity (ΔF/F) was calculated using the equation: ΔF/F = (F-F_0_)/F_0_, where F_0_ is defined as the average fluorescence intensity of the consecutive 5 time points before spine stimulation, and F is defined as the fluorescence intensity measured at the time point of interest.

### Ex vivo ER-HaloCaMP Ca^2+^ imaging in the Drosophila brain

Drosophila brain live imaging experiments were carried out as in Silva et al.^92^ with some modifications. Briefly, female flies carrying the Myosuppressin (Ms)-Gal4 driver9^3^ were crossed with male flies carrying the UAS-ER-HaloCaMP-BFP construct (which is fused to mTagBFP2, as in the case of experiments in cells in culture). Crosses for imaging experiments were raised at 25°C. 1-to 2-day-old adult progeny were used for each recording.

A single fly brain was dissected in haemolymph solution and mounted on a glass coverslip coated with poly-L-lysine (Sigma-Aldrich, P1524). The haemolymph solution contained 130mM NaCl (Sigma, S9625), 5 mM KCl (Sigma, P3911), 2 mM MgC1_2_ (Sigma, M9272), 2 mM CaC1_2_ (Sigma, C3881), 5 mM D-trehalose (Sigma, T9531), 30 mM sucrose (Sigma, S9378) and 5 mM HEPES-hemisodium salt (Sigma, H7637). Then, a total volume of 500 µL ofhaemolymph solution was added on top of the coverslip. Next, 1 µL of the JF_58_rHTL 500 µM stock solution was incubated for approximately 40 min prior imaging.

Two-photon imaging was performed on a Leica Stellaris 8 DIVE upright microscope equipped with a × 25, 1.00-NA water-immersion objective. Two-photon excitation of mTagBFP2 and ER-HaloCaMP_585_ was achieved using a Mai Tai eHP DeepSee MP laser tuned to 820 nm. Spectral separation was achieved by tuning detectors for mTagBFP2 (445-460nm) and JF_585_-HTL (580-650nm). Then, 512 × 512 images were acquired at 0.5Hz, and the entire duration of each recording was 1200 s. For Ca^2+^ depletion experiments, Thapsigargin (Sigma, T9033) was diluted in haemolymph solution to prepare a 100 µM stock solution. After 60 s of baseline acquisition, 5 µ1of the Thapsigargin stock solution was added to the 500-µl bath solution on top of the brain, resulting in a final concentration of 1 µM.

Image analysis was performed using a custom-written MATLAB script^92^. Regions of interest (ROis) were manually drawn around individual Myosuppressin neuronal somas. The average intensity values for mTagBFP2 and ER-HaloCaMP_585_ channels over each ROI were calculated over time after background subtraction. The ratio was calculated by dividing ER-HaloCaMP_585_ by mTagBFP2 intensity. Traces from all cell bodies were pooled for analysis.

### Image analysis and statistics

We used the ImageJ plugin Time Series Analyzer V3 for imaging analysis of dynamic changes in fluorescence, except for ex vivo experiments in which we used a custom-written MATLAB script^92^. Statistical analysis was carried out using GraphPad Prism v10 for Windows, with specific tests indicated in the Supplementary Table STl. For each dataset, normality was assessed using the Shapiro-Wilk test and then based on the result we chose either parametric or non-parametric tests for further comparisons. All n numbers, as well as the number of independent experiments, are detailed in the Supplementary Table STl. Unless otherwise indicated, all data for this study was acquired from at least 3 independent experiments. We used Python to calculate Spearman correlations and their associated p-values with the SciPy library. Pearson’s correlation coefficients and associated p values were calculated using graphpad, and both were incorporated in a single heatmap using illustrator. For the figures, the mean and the standard error of mean are used. Results of statistical analysis are shown in figures corresponding to the following criteria: *p<0.05, **p<0.01, ***p<0.001, ****p < 0.0001, n.s., not statistically significant.

## Author contributions

Conceptualization, J.d.J-S.; methodology, A.M., H.F., R.F., K.Z., B.S, C.R., J.d.J.S.; investigation, A.M., H.F., R.F., V. R., B.S., C.R., K.Z., J.d.J.S.; project administration and supervision, J.d.J-S.; writing - original draft, A.M. and J.d.J-S.; writing - review & editing. A.M., H.F., R.F., K.Z., B.S., C.R., C.G., T.O., D.H., V.R., E. R .S. and J.d.J.S.

## Acknowledgements

We would like to thank all members of the de Juan-Sanz laboratory for insightful discussions and comments. We thank Dr. Luke Lavis and the Janelia Open Science Initiative for providing the Janelia Fluor dyes used in this study. Part of this work was carried out in the ICM.Quant core facility (RRID:SCR_026393). We gratefully acknowledge Claire Lovo, Meryem Rezzik and Astou Tangara for setting up multiplexing experiments. Animals were hosted with the help of the PHENOPARC facility at ICM.This work was made possible by the Paris Brain Institute Diane Barriere Chair in Synaptic Bioenergetics awarded to J.d.J.-S., funding from the French national program ‘‘Investissements d’avenir’’ ANR-10-IAIHU-0006 awarded to the ICM and funding by the Richard Mille Fund (Project DBS, 2023-2028). Our additional funding sources are an ERC Starting Grant SynaptoEnergy (European Research Council, ERC-StG-852873), 2019 ATIP-Avenir Grant (CNRS, lnserm), and two grants by the French National Research Agency (ANR) under project numbers ANR-24-CE16-0221 and ANR-22-CE16-0020 awarded to J.d.J.-S. D.H. is funded by an ERC Starting Grant GutSense (European Research Council, ERC-StG-101117267). A.M. is the recipient of a predoctoral fellowship from the French Ministry of Science. J.d.J.-S. is a permanent CNRS researcher and a FENS-Kavli Scholar. V.R. is supported by the Max Planck Society.

## Declaration of interests

The authors declare no competing interests.

**Fig. S1.**
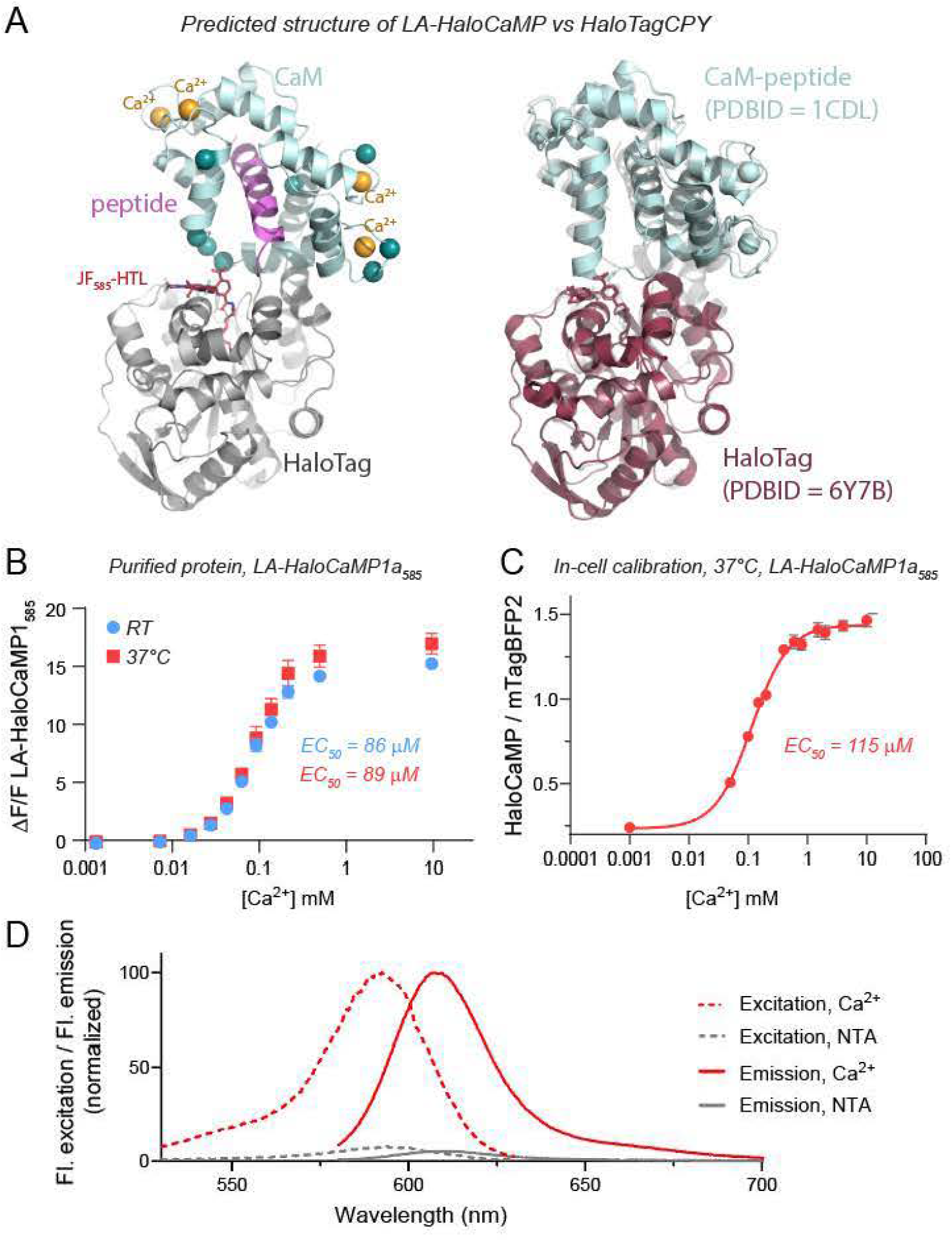
Biochemical properties of LA-HaloCaMPl. **(A)** Predicted structural models of low-affinity HaloCaMP (LA-HaloCaMP) and HaloTagCPY generated using Chai Discovery. Left: LA-HaloCaMP is shown with HaloTag (gray), the CaM-binding peptide (purple), calmodulin (CaM, cyan) and bound calcium ions (yellow). JF_585_,HTL is represented in red Right: HaloTagCPY is modeled as a fusion ofHaloTag (PDB ID: 6Y7B) and the CaM--peptide complex (PDB ID: lCDL). **(B)** In vitro calcium titration of purified LA-HaloCaMP_585_ protein at room temperature (RT, blue) and 37°C (red), showing EC_50_ of 86 µM at room temperature (RT) and 89 µMat 37°C. (C) *In-cell* calibration at 37°C of LA-HaloCaMP_5_ss in permeabilized HeLa cells, yielding an EC_50_ of 115 µM. (D) Excitation and emission spectra of LA-HaloCaMP labeled with JF_585-_HTL in the presence of Ca^2+^ (red) or Ca^2+^-free buffered with NTA (gray). Excitation spectra (dashed lines) and emission spectra (solid lines) were normalized to their respective maxima. Data are represented as mean ± SEM. See Supplementary Table STl for details on sample sizes.

**Fig. S2.**
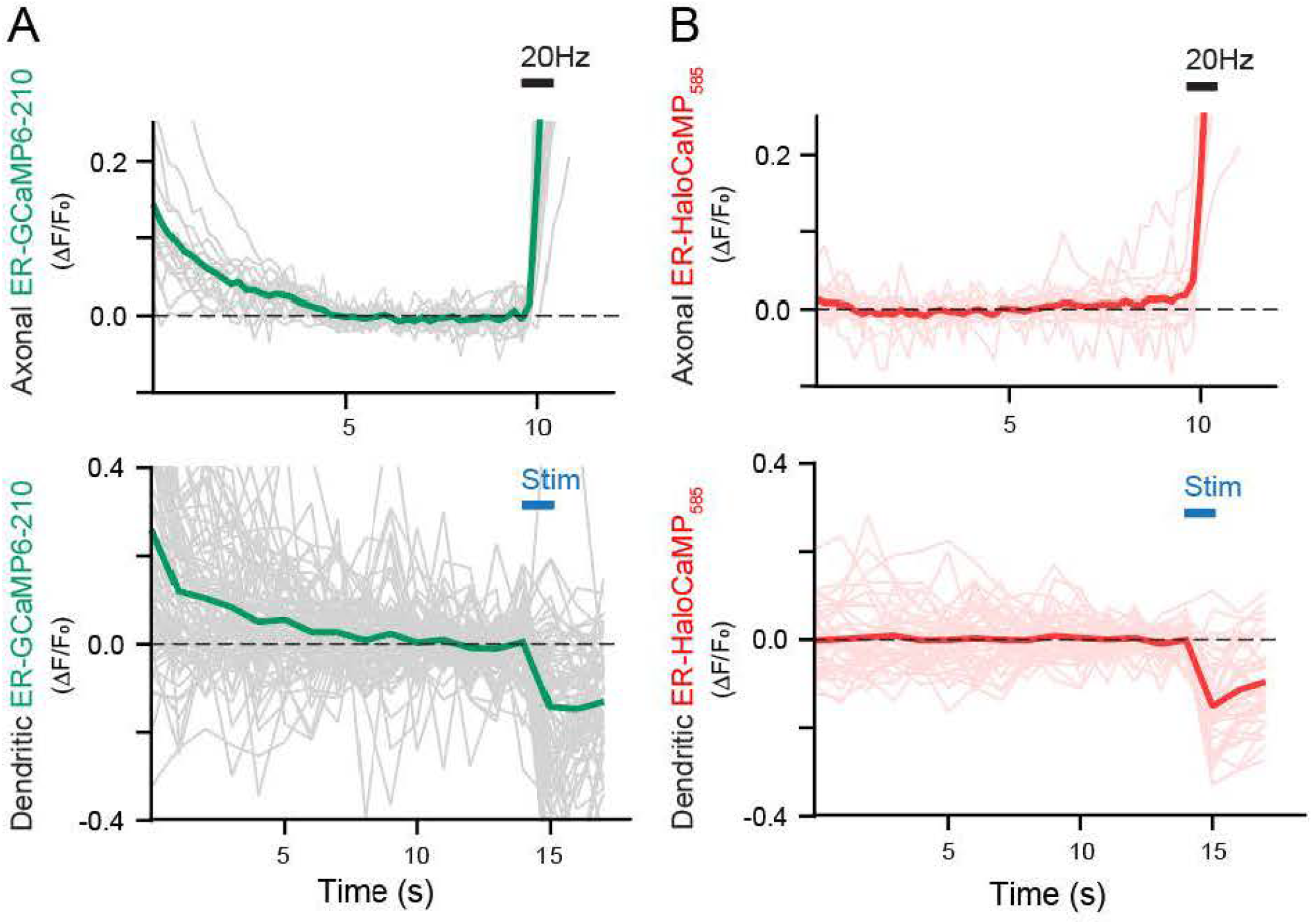
Photostability comparisons in axons and dendrites after light exposure. **(A)** Imaging of ER-GCaMP6-210 in either axons (top) or dendrites (bottom) shows a quick lossin fluorescence compatible with photoswitching of a fraction of the sensor population, resulting in a first decay in fluorescence that is quickly stabilized. **(B)** Comparative measurements of ER-HaloCaMPsis_585_ in axons (top) or dendrites (bottom) show that signals were stable right after the illumination of the sample starts, showing improved photostability. Data are represented as mean ± SEM. See Supplementary Table STl for details on sample sizes.

**Fig. S3.**
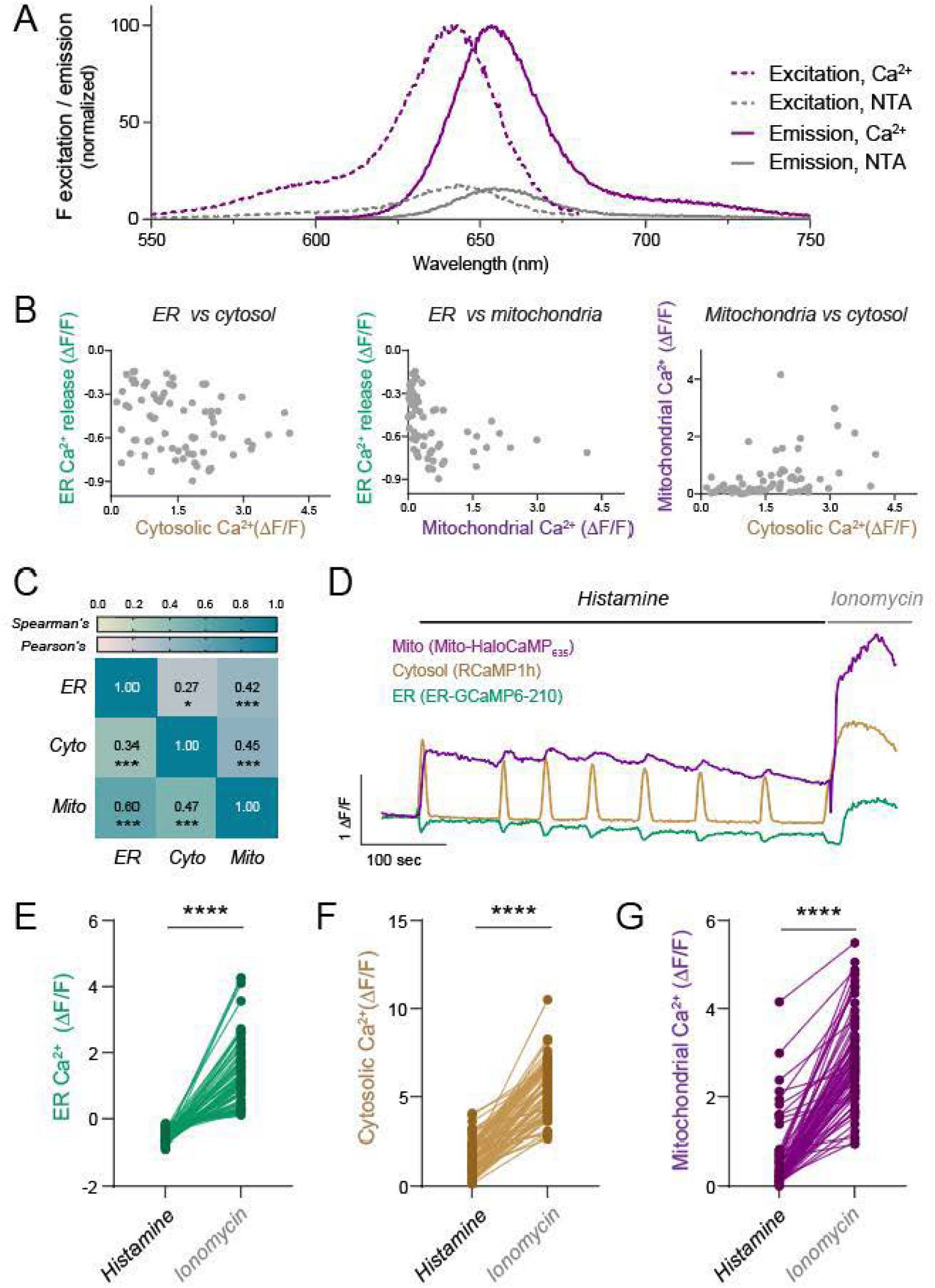
Correlation analysis of Cai+ retpolllM and non-uturatlon of die sensors during histamine. **(A)** Single-cell Ca^2+^ peak responses to histamine in theERand cytosol (left), ER and mitochondria (middle), or mitochondria and cy10S01 (right). **(B)** Correlation matrix of the data shown in (A), calculated under two assumptions: linear relationships (Pearson’s) and mono10nic relationships (Spearman’s). Each cell shows the respective correlation coefficient and asterisks indicate statistically significant deviation from zero. **(C)** Paired single-cell Ca^2+^ peak responses in the ER (left), cytosol (middle), and mitochondria (right) after histamine and subsequent ionomycin treatment. Ionomycin saturates each sensor and provides a signal that is always higher than the histamine peak response, confirming that none of the sensors was saturated by histamine. See Supplementary Table STl for details on statistical tests and sample sizes.

**Fig. S4.**
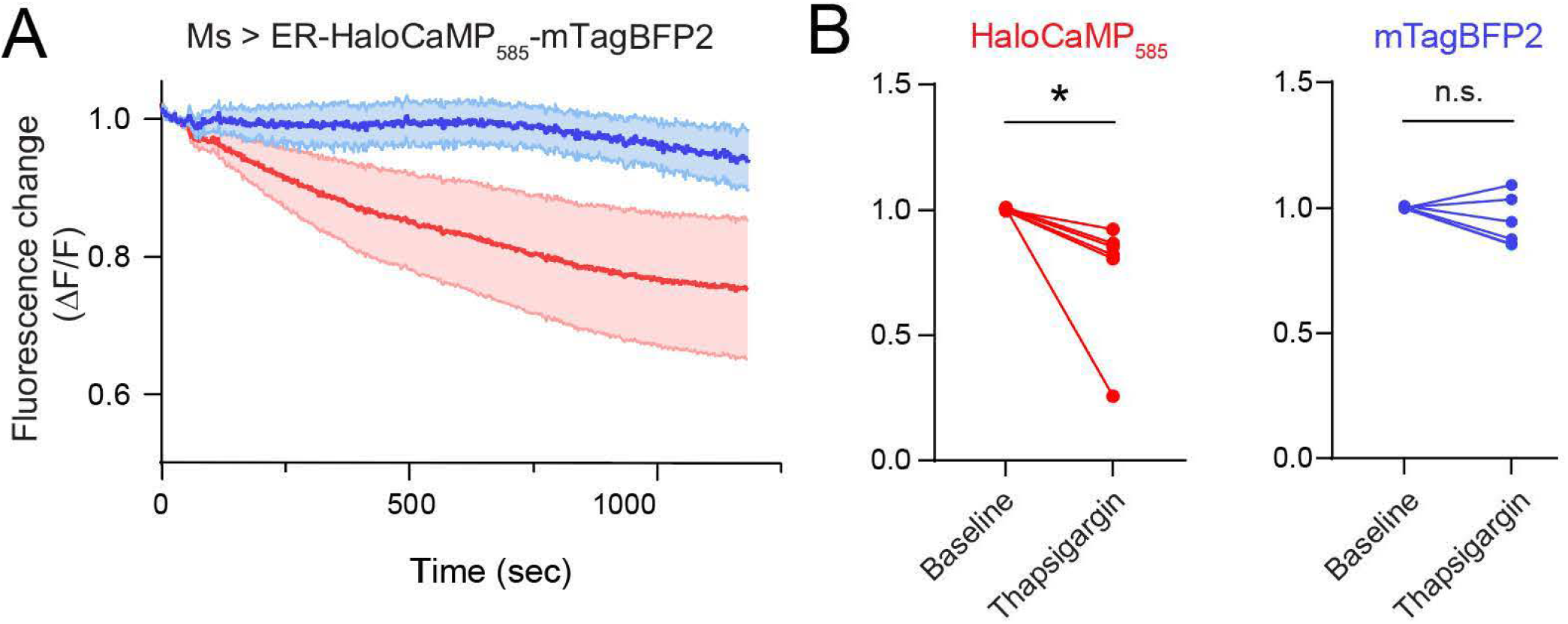
Simultaneous measurements of ER-HaloCaMP_585_ and mTagBFP2 in Ms fly neurons expressing ER-HaloCaMP_585_. **(A)** Mean relative fluorescence changes normalized to baseline over time in response to ER calcium depletion induced by thapsigargin in LA-HaloCaMP_585_ and mTagBFP2. Shaded areas represent SEM. **(B)** Quantification of fluorescence changes for ER-HaloCaMP585 (left, red) and mTagBFP2 (right, blue) at baseline and after thapsigargin application in paired samples. ER-HaloCaMP_585_ fluorescence significantly decreased whereas mTagBFP2 fluorescence remained stable (n.s., not significant). See Supplementary Table STl for details on statistical tests and sample sizes

## References

1. Berridge, M.J., Lipp, P., and Bootman, M.D. (2000). The versatility and universality of calcium signalling. Nature reviews. Molecular cell biology 1, 11–21. 10.1038/35036035.

2. Raffaello, A., Mammucari, C., Gherardi, G., and Rizzuto, R. (2016). Calcium at the center of cell signaling: interplay between endoplasmic reticulum, mitochondria and lysosomes. Trends in biochemical sciences 41, 1035. 10.1016/J.TIBS.2016.09.001.

3. Mekahli, D., Bultynck, G., Parys, J.B., De Smedt, H., and Missiaen, L. (2011). Endoplasmic-reticulum calcium depletion and disease. Cold Spring Harbor perspectives in biology 3, a004317.. 10.1101/cshperspect.a004317.

4. Rizzuto, R., De Stefani, D., Raffaello, A., and Mammucari, C. (2012). Mitochondria as sensors and regulators of calcium signalling. Nature reviews. Molecular cell biology 13, 566---578. 10.1038/nrm3412.

5. Dematteis, G., Tapella, L., Casali, C., Talmon, M., Tonelli, E., Reano, S., Ariotti, A., Pessolano, E., Malecka, J., Chrostek, G., et al. (2024). ER-mitochondria distance is a critical parameter for efficient mitochondrial Ca2+ uptake and oxidative metabolism. Commun Biol 7, 1-15. 10.1038/s42003-024-06933-9.

6. Chakrabarti, R., Ji, W.-K., Stan, R.V., de Juan Sanz, J., Ryan, T.A., and Higgs, H.N. (2018). INF2-mediated actin polymerization at the ER stimulates mitochondrial calcium uptake, inner membrane constriction, and division. The Journal of Cell Biology 217, 251–268. 10.1083/jcb.201709111.

7. Divakaruni, S.S., Dyke, A.M.V., Chandra, R., LeGates, T.A., Contreras, M., Dharmasri, P.A., Higgs, H.N., Lobo, M.K., Thompson, S.M., and Blanpied, T.A. (2018). Long-term potentiation requires a rapid burst of dendritic mitochondrial fission during induction. Neuron 100, 860. 10.1016/J.NEURON.2018.09.025.

8. Cardenas, C., and Foskett, J.K. (2012). Mitochondrial Ca2+ signals in autophagy. Cell Calcium 52,44--51. 10.1016/j.ceca.2012.03.001.

9. Park, S.J., Lee, S.B., Suh, Y., Kim, S.-J., Lee, N., Hong, J.-H., Park, C., Woo, Y., Ishizuka, K., Kim, J.-H., et al. (2017). DISCl Modulates Neuronal Stress Responses by Gate-Keeping ER-Mitochondria Ca2+ Transfer through the MAM. Cell reports 21, 2748–2759. 10.1016/j.celrep.2017.11.043.

10. Panzera, L.C., Johnson, B., Quinn, J.A., Cho, I.H., Tamkun, M.M., and Hoppa, M.B. (2022). Activity-dependent endoplasmic reticulum Ca2+ uptake depends on Kv2.1-mediated endoplasmic reticulum/plasma membrane junctions to promote synaptic transmission. Proceedings of the National Academy of Sciences 119, e2117135119. 10.1073/pnas.2117135119.

11. Ashrafi, G., de Juan-Sanz, J., Farrell, R.J., and Ryan, T.A. (2020). Molecular Tuning of the Axonal Mitochondrial Ca2+ Uniporter Ensures Metabolic Flexibility ofNeurotransmission. Neuron 105, 678-687.e5.10.1016/j.neuron.2019.11.020.

12. de Juan-Sanz, J., Holt, G.T., Schreiter, E.R., de Juan, F., Kirn, D.S., and Ryan, T.A. (2017). Axonal Endoplasmic Reticulum Ca2+ Content Controls Release Probability in CNS Nerve Terminals. Neuron 93, 867-881.e6. 10.1016/j.neuron.2017.01.010.

13. Vaccaro, V., Devine, M.J., Higgs, N.F., and Kittler, J.T. (2017). Mirol-dependent mitochondrial positioning drives the rescaling of presynaptic Ca2+ signals during homeostatic plasticity. EMBO reports 18, 231–240. 10.15252/EMBR.201642710.

14. Kwon, S.K., Sando, R., Lewis, T.L., Hirabayashi, Y., Maximov, A., and Polleux, F. (2016). LKBl Regulates Mitochondria-DependentPresynaptic Calcium Clearance and Neurotransmitter Release Properties at Excitatory Synapses along Cortical Axons. PLOS Biology 14, e1002516. 10.1371/JOURNAL.PBIO.1002516.

15. Liiv, M., Vaarmann, A., Safiulina, D., Choubey, V., Gupta, R., Kuum, M., Janickova, L., Hodurova, Z., Cagalinec, M., Zeb, A., et al. (2024). ER calcium depletion as a key driver for impaired ER-to-mitochondria calcium transfer and mitochondrial dysfunction in Wolfram syndrome. Nat Commun 15, 6143. 10.1038/s41467-024-50502-x.

16. Lee, K.-S., Huh, S., Lee, S., Wu, Z., Kim, A.-K., Kang, H.-Y., and Lu, B. (2018). Altered ER-mitochondria contact impacts mitochondria calcium homeostasis and contributes to neurodegeneration in vivo in disease models. Proc Natl Acad Sci U SA 115, E8844--E8853. 10.1073/pnas.1721136115.

17. de Ridder, I., Kerkhofs, M., Lemos, F.O., Loncke, J., Bultynck, G., and Parys, J.B. (2023). The ER-mitochondria interface, where Ca2+ and cell death meet. Cell Calcium 112, 102743. 10.1016/j.ceca.2023.102743.

18. Loncke, J., Kaasik, A., Bezprozvanny, I., Parys, J.B., Kerkhofs, M., and Bultynck, G. (2021). Balancing ER-Mitochondrial Ca2+ Fluxes in Health and Disease. Trends in Cell Biology 31, 598–612. 10.1016/j.tcb.2021.02.003.

19. Suzuki, J., Kanemaru, K., Ishii, K., Ohkura, M., Okubo, Y., and Iino, M. (2014). Imaging intraorganellar Ca2+ at subcellular resolution using CEPIA. Nature communications 5, 4153. 10.1038/ncomms5153.

20. Bovo, E., Martin, J.L., Tyryfter, J., de Tombe, P.P., and Zima, A.V. (2016). R-CEPIAler as a new tool to directly measure sarcoplasmic reticulum [Ca] in ventricular myocytes. Am J Physiol Heart Circ Physiol 311, H268---H275. 10.1152/ajpheart.00175.2016.

21. Wu, J., Prole, D.L., Shen, Y., Lin, Z., Gnanasekaran, A., Liu, Y., Chen, L., Zhou, H., Chen, S.R.W., Usachev, Y.M., et al. (2014). Red fluorescent genetically encoded Ca 2+ indicators for use in mitochondria and endoplasmic reticulum. Biochemical Journal 464, 13–22. 10.1042/BJ20140931.

22. Henderson, M.J., Baldwin, H.A., Werley, C.A., Boccardo, S., Whitaker, L.R., Yan, X., Holt, G.T., Schreiter, E.R., Looger, L.L., Cohen, A.E., et al. (2015). A Low Affinity GCaMP3 Variant (GCaMPer) for Imaging the Endoplasmic Reticulum Calcium Store. PloS one 10, e0139273. 10.1371/journal.pone.0139273.

23. Okubo, Y., Suzuki, J., Kanemaru, K., Nakamura, N., Shibata, T., and Iino, M. (2015). Visualization of Ca2+ Filling Mechanisms upon Synaptic Inputs in the Endoplasmic Reticulum of Cerebellar Purkinje Cells. The Journal of neuroscience : the official journal of the Society for Neuroscience 35, 15837–15846. 10.1523/JNEUROSCI.3487-15.2015.

24. Hirabayashi, Y., Kwon, S.K., Paek, H., Pernice, W.M., Paul, M.A., Lee, J., Erfani, P., Raczkowski, A., Petrey, D.S., Pon, L.A., et al. (2017). ER-mitochondria tethering by PDZD8 regulates Ca2+ dynamics in mammalian neurons. Science 358, 623–630. 10.1126/science.aan6009.

25. Wang, M., Da, Y., and Tian, Y. (2023). Fluorescent proteins and genetically encoded biosensors. Chem. Soc. Rev. 52, 1189–1214. 10.1039/D2CS00419D.

26. Deisseroth, K., and Hegemann, P. (2017). The form and function of channelrhodopsin. Science 357, eaan5544. 10.l126/science.aan5544.

27. Chen, Z., Truong, T.M., and Ai, H. (2017). Illuminating Brain Activities with Fluorescent Protein-Based Biosensors. Chemosensors 5, 32. 10.3390/chemosensors5040032.

28. Shaner, N.C., Steinbach, P.A., and Tsien, R.Y. (2005). A guide to choosing fluorescent proteins. Nat Methods 2, 905–909. 10.1038/nmeth819.

29. lcha, J., Weber, M., Waters, J.C., and Norden, C. (2017). Phototoxicity in live fluorescence microscopy, and how to avoid it. Bioessays 39. 10.1002/bies.201700003.

30. Laissue, P.P., Alghamdi, R.A., Tomancak, P., Reynaud, E.G., and Shroff, H. (2017). Assessing phototoxicity in live fluorescence imaging. Nat Methods 14, 657–661. 10.1038/nmeth.4344.

31. Los, G.V., Encell, L.P., McDougall, M.G., Hartzell, D.D., Karassina, N., Zimprich, C., Wood, M.G., Learish, R., Ohana, R.F., Urh, M., et al. (2008). HaloTag: A Novel Protein Labeling Technology for Cell Imaging and Protein Analysis. ACS Chem. Biol. 3, 373–382. 10.1021/cb800025k.

32. Abdelfattah, A.S., Kawashima, T., Singh, A., Novak, O., Liu, H., Shuai, Y., Huang, Y.-C., Campagnola, L., Seeman, S.C., Yu, J., et al. (2019). Bright and photostable chemigenetic indicators for extended in vivo voltage imaging. Science 365, 699–704. 10.1126/science.aav6416.

33. Abdelfattah, A.S., Valenti, R., Zheng, J., Wong, A., GENIE Project Team, Podgorski, K., Koyama, M., Kim, D.S., and Schreiter, E.R. (2020). A general approach to engineer positive-going eFRET voltage indicators. Nat Commun 11, 3444. 10.1038/s41467-020-l7322-l.

34. Hellweg, L., Edenhofer, A., Barck, L., Huppertz, M.-C., Frei, M.S., Tamawski, M., Bergner, A., Koch, B., Johnsson, K., and Hiblot, J. (2023). A general method for the development of multicolorbiosensors with large dynamic ranges. Nat Chem Biol 19, 1147–1157. 10.1038/s41589-023-01350-1.

35. Zheng, Y., Cai, R., Wang, K., Zhang, J., Zhuo, Y., Dong, H., Zhang, Y., Wang, Y., Deng, F., Ji, E., et al. (2024). In vivo multiplex imaging of dynamic neurochemical networks with designed far-red dopamine sensors. Preprint at bioRxiv, 10.1101/2024.12.22.629999 https://doi.org/10.1101/2024.12.22.629999.

36. Asanuma, D., Takaoka, Y., Namiki, S., Takikawa, K., Kamiya, M., Nagano, T., Urano, Y., and Hirose, K. (2014). Acidic-pH-activatable fluorescence probes for visualizing exocytosis dynamics. Angew Chem Int Ed Engl 53, 6085–6089. 10.1002/anie.201402030.

37. Liu, Y., Miao, K., Li, Y., Fares, M., Chen, S., and Zhang, X. (2018). A HaloTag-Based Multicolor Fluorogenic Sensor Visualizes and Quantifies Proteome Stress in Live Cells Using Solvatochromic and Molecular Rotor-Based Fluorophores. Biochemistry 57, 4663–4674. 10.1021/acs.biochem.8b00135.

38. Cook, A., Walterspiel, F., and Deo, C. (2023). HaloTag-Based Reporters for Fluorescence Imaging and Biosensing. ChemBioChem 24, e202300022. 10.1002/cbic.202300022.

39. Farrants, H., Hiblot, J., Griss, R., and Johnsson, K. (2017). Rational Design and Applications of Semisynthetic Modular Biosensors: SNIFITs and LUCIDs. Methods Mol Biol 1596, 101-117. 10.1007/978-1-4939-6940-1_7.

40. Deo, C., Abdelfattah, A.S., Bhargava, H.K., Berro, A.J., Falco, N., Farrants, H., Moeyaert, B., Chupanova, M., Lavis, L.D., and Schreiter, E.R. (2021). The HaloTag as a general scaffold for far-red tunable chemigenetic indicators. Nature Chemical Biology 17. 10.1038/s41589-021-00775-w.

41. Farrants, H., Shuai, Y., Lemon, W.C., Monroy Hernandez, C., Zhang, D., Yang, S., Patel, R., Qiao, G., Frei, M.S., Plutkis, S.E., et al. (2024). A modular chemigenetic calcium indicator for multiplexed in vivo functional imaging. Nat Methods 21, 1916–1925. 10.1038/s41592-024-02411-6.

42. Deo, C., Sheu, S.-H., Seo, J., Clapham, D.E., and Lavis, L.D. (2019). Isomeric Tuning Yields Bright and Targetable Red Ca2+ Indicators. J Am Chem Soc 141, 13734--13738. 10.1021/jacs.9b06092.

43. Mertes, N., Busch, M., Huppertz, M.-C., Hacker, C.N., Wilhelm, J., Giirth, C.-M., Kuhn, S., Hiblot, J., Koch, B., and Johnsson, K. (2022). Fluorescent and Bioluminescent Calcium Indicators with Tuneable Colors and Affinities. J Am Chem Soc 144, 6928–6935. 10.1021/jacs.2c01465.

44. Zhao, Y., Araki, S., Wu, J., Teramoto, T., Chang, Y.-F., Nakano, M., Abdelfattah, A.S., Fujiwara, M., Ishihara, T., Nagai, T., et al. (2011). An Expanded Palette of Genetically Encoded Ca2+ Indicators. Science 333, 1888---1891. 10.l126/science.1208592.

45. Dana, H., Mohar, B., Sun, Y., Narayan, S., Gordus, A., Hasseman, J.P., Tsegaye, G., Holt, G.T., Hu, A., Walpita, D., et al. (2016). Sensitive red protein calcium indicators for imaging neural activity. eLife 5. 10.7554/eLife.12727.

46. Inoue, M., Takeuchi, A., Horigane, S., Ohkura, M., Gengyo-Ando, K., Fujii, H., Kamijo, S., Takemoto-Kimura, S., Kano, M., Nakai, J., et al. (2015). Rational design of a high-affinity, fast, red calcium indicatorR-CaMP2. Nat Methods 12, 64–70. 10.1038/nmeth.3185.

47. Inoue, M., Takeuchi, A., Manita, S., Horigane, S., Sakamoto, M., Kawakami, R., Yamaguchi, K., Otomo, K., Yokoyama, H., Kim, R., et al. (2019). Rational Engineering of XCaMPs, a Multicolor GECI Suite for In Vivo Imaging of Complex Brain Circuit Dynamics. Cell 177, 1346-1360.e24. 10.1016/j.cell.2019.04.007.

48. Yokoyama, T., Manita, S., Uwamori, H., Tajiri, M., Imayoshi, I., Yagishita, S., Murayama, M., Kitamura, K., and Sakamoto, M. (2024). A multicolor suite for deciphering population coding of calcium and cAMP in vivo. Nat Methods 21, 897–907. 10.1038/s41592-024-02222-9.

49. Fink, R., Imai, S., Gockel, N., Lauer, G., Renken, K., Wietek, J., Lamothe-Molina, P.J., Fuhrmann, F., Mittag, M., Ziebarth, T., et al. (2025). PinkyCaMP a mScarlet-based calcium sensor with exceptional brightness, photostability, and multiplexing capabilities. Preprint at bioRxiv, 10.1101/2024.12.16.628673 https://doi.org/10.1101/2024.12.16.628673.

50. Farrell, R.J., Bredvik, K.G., Hoppa, M.B., Hennigan, S.T., Brown, T.A., and Ryan, T.A. (2024). A ratiometric ER calcium sensor for quantitative comparisons across cell types and subcellular regions. bioRxiv, 02.15.580492. 10.1101/2024.02.15.580492.

51. Thastrup, O., Cullen, P.J., Drnbak, B.K., Hanley, M.R., and Dawson, A.P. (1990). Thapsigargin, a tumor promoter, discharges intracellular Ca2+ stores by specific inhibition of the endoplasmic reticulum Ca2(+)-ATPase. Proceedings of the National Academy of Sciences 87, 2466--2470. 10.1073/pnas.87.7.2466.

52. Courjaret, R., Dib, M., and Machaca, K. (2018). Spatially restricted subcellular Ca2+ signaling downstream of store-operated calcium entry encoded by a cortical tunneling mechanism. Sci Rep 8, 11214. 10.1038/s41598-018-29562-9.

53. Terasaki, M. (2018). Axonal endoplasmic reticulum is very narrow. Journal of cell science 131, jcs210450. 10.1242/jcs.210450.

54. Wu, Y., Whiteus, C., Xu, C.S., Hayworth, K.J., Weinberg, R.J., Hess, H.F., and De Camilli, P. (2017). Contacts between the endoplasmic reticulum and other membranes in neurons. Proceedings of the National Academy of Sciences of the United States of America 114, E4859---E4867. 10.1073/pnas.1701078114.

55. Spacek, J., and Harris, K.M. (1997). Three-Dimensional Organization of Smooth Endoplasmic Reticulum in Hippocampal CAI Dendrites and Dendritic Spines of the Immature and Mature Rat. J. Neurosci. 17, 190–203. 10.1523/JNEUROSCI.17-01-00190.1997.

56. Oliva, M.K., Perez-Moreno, J.J., O’Shaughnessy, J., Wardill, T.J., and O’Kane, C.J. (2020). Endoplasmic Reticulum Lumenal Indicators in Drosophila Reveal Effects ofHSP-Related Mutations on Endoplasmic Reticulum Calcium Dynamics. Frontiers in Neuroscience 14, 816. 10.3389/fnins.2020.00816.

57. Grieves, R.M., Wood, E.R., and Dudchenko, P.A. (2016). Place cells on a maze encode routes rather than destinations. eLife 5. 10.7554/ELIFE.15986.

58. Benedetti, L., Fan, R., Weigel, A.V., Moore, A.S., Houlihan, P.R., Kittisopikul, M., Park, G., Petruncio, A., Hubbard, P.M., Pang, S., et al. (2025). Periodic ER-plasma membrane junctions support long-range Ca2+ signal integration in dendrites. Cell 188, 484-500.e22. 10.1016/j.cell.2024.11.029.

59. Holbro, N., Grunditz, A., and Gertner, T.G. (2009). Differential distribution of endoplasmic reticulum controls metabotropic signaling and plasticity at hippocampal synapses. Proceedings of the National Academy of Sciences 106, 15055–15060. 10.1073/pnas.0905110106.

60. Bapat, O., Purimetla, T., Kruessel, S., Shah, M., Fan, R., Thum, C., Rupprecht, F., Langer, J.D., and Rangaraju, V. (2024). VAP spatially stabilizes dendritic mitochondria to locally support synaptic plasticity. Nat Commun 15, 205. 10.1038/s41467-023-44233-8.

61. Jain, A., Nakahata, Y., Pancani, T., Watabe, T., Rusina, P., South, K., Adachi, K., Yan, L., Simorowski, N., Furukawa, H., et al. (2024). Dendritic, delayed, stochastic CaMKII activation in behavioural time scale plasticity. Nature 635, 151–159. 10.1038/s41586-024-08021-8.

62. Andresen, M., Stiel, A.C., Trowitzsch, S., Weber, G., Eggeling, C., Wahl, M.C., Hell, S.W., and Jakobs, S. (2007). Structural basis for reversible photoswitching in Dronpa. Proceedings of the National Academy of Sciences 104, 13005–13009. 10.1073/pnas.0700629104.

63. Zhou, Z., Matlib, M.A., and Bers, D.M. (1998). Cytosolic and mitochondrial Ca2+ signals in patch clamped mammalian ventricular myocytes. J Physiol 507 (Pt 2), 379–403. 10.1111/j.1469-7793.1998.379bt.x.

64. Di Lisa, F., Gambassi, G., Spurgeon, H., and Hansford, R.G. (1993). Intramitochondrial free calcium in cardiac myocytes in relation to dehydrogenase activation. Cardiovasc Res 27, 1840-1844. 10.1093/cvr/27.10.1840.

65. Miyata, H., Silverman, H.S., Sollott, S.J., Lakatta, E.G., Stem, M.D., and Hansford, R.G. (1991). Measurement of mitochondrial free Ca2+ concentration in living single rat cardiac myocytes. Am J Physiol 261, Hl 123–1134. 10.l152/ajpheart.1991.261.4.Hl123.

66. Groten, C.J., and MacVicar, B.A. (2022). Mitochondrial Ca2+ uptake by the MCU facilitates pyramidal neuron excitability and metabolism during action potential firing. Commun Biol 5, 1–15. 10.1038/s42003-022-03848-1.

67. Bredvik, K., and Ryan, T.A. (2024). Differential Control of Inhibitory and Excitatory Nerve Terminal Function by Mitochondria. Preprint at bioRxiv, 10.1101/2024.05.19.594864 https://doi.org/10.1101/2024.05.19.594864.

68. Chen-Engerer, H.-J., Hartmann, J., Karl, R.M., Yang, J., Feske, S., and Konnerth, A. (2019). Two types of functionally distinct Ca2+ stores in hippocampal neurons. Nat Commun 10, 3223. 10.1038/s41467-019-11207-8.

69. Hartmann, J., Karl, R.M., Alexander, R.P.D., Adelsberger, H., Brill, M.S., Riihlmann, C., Ansel, A., Sakimura, K., Baba, Y., Kurosaki, T., et al. (2014). STIMl controls neuronal Ca2+ signaling, mGluRl-dependent synaptic transmission, and cerebellar motor behavior. Neuron 82, 635–644. 10.1016/j.neuron.2014.03.027.

70. Kar, P., Samanta, K., and Parekh, A.B. (2020). Cytosolic and intra-organellar Ca2+ oscillations: mechanisms and function. Current Opinion in Physiology 17, 175–186. 10.1016/j.cophys.2020.08.011.

71. Arruda, A.P., and Hotamisligil, G.S. (2015). Calcium homeostasis and organelle function in the pathogenesis of obesity and diabetes. Cell Metab 22, 381–397. 10.1016/j.cmet.2015.06.010.

72. Zheng, S., Wang, X., Zhao, D., Liu, H., and Hu, Y. (2023). Calcium homeostasis and cancer: insights from endoplasmic reticulum-centered organelle communications. Trends in Cell Biology 33, 312–323. 10.1016/j.tcb.2022.07.004.

73. Liu, Y., Ma, X., Fujioka, H., Liu, J., Chen, S., and Zhu, X. (2019). DJ-1 regulates the integrity and function of ER-mitochondria association through interaction with IP3R3-Grp75-VDAC1. Proceedings of the National Academy of Sciences 116, 25322–25328. 10.1073/pnas.1906565116.

74. Paillusson, S., Stoica, R., Gomez-Suaga, P., Lau, D.H.W., Mueller, S., Miller, T., and Miller, C.C.J. (2016). There’s Something Wrong with my MAM; the ER-Mitochondria Axis and Neurodegenerative Diseases. Trends in Neurosciences 39, 146--157. 10.1016/j.tins.2016.01.008.

75. Stoica, R., De Vos, K.J., Paillusson, S., Mueller, S., Sancho, R.M., Lau, K.-F., Vizcay-Barrena, G., Lin, W.-L., Xu, Y.-F., Lewis, J., et al. (2014). ER-mitochondria associations are regulated by the VAPB-PTPIP51 interaction and are disrupted by ALS/FTD-associated TDP-43. Nat Commun 5, 3996. 10.1038/ncomrns4996.

76. Si, H., Wang, J., Meininger, C.J., Peng, X., Zawieja, D.C., and Zhang, S.L. (2020). Ca2+ release-activated Ca2+ channels are responsible for histamine-induced Ca2+ entry, permeability increase, and interleukin synthesis in lymphatic endothelial cells. American Journal of Physiology-Heart and Circulatory Physiology 318, H1283–H1295. 10.1152/ajpheart.00544.2019.

77. Cartes-Saavedra, B., Macuada, J., Lagos, D., Arancibia, D., Andres, M.E., Yu-Wai-Man, P., Hajn6czky, G., and Eisner, V. (2022). OPAi Modulates Mitochondrial Ca2+ Uptake Through ER-Mitochondria Coupling. Front Cell Dev Biol 9, 774108. 10.3389/fcell.2021.774108.

78. Lopez-Crisosto, C., Diaz-Vegas, A., Castro, P.F., Rothermel, B.A., Bravo-Sagua, R., and Lavandero, S. (2021). Endoplasmic reticulum-mitochondria coupling increases during doxycycline-induced mitochondrial stress in HeLa cells. Cell Death Dis 12, 1–12. 10.1038/s41419-021-03945-9.

79. de Brito, O.M., and Scorrano, L. (2008). Mitofusin 2 tethers endoplasmic reticulum to mitochondria. Nature 456, 605---610. 10.1038/nature07534.

80. Filadi, R., Greotti, E., Turacchio, G., Luini, A., Pozzan, T., and Pizzo, P. (2015). Mitofusin 2 ablation increases endoplasmic reticulum-mitochondria coupling. Proc Natl Acad Sci US A 112, E2174–E2181. 10.1073/pnas.1504880112.

81. Patron, M., Granatiero, V., Espino, J., Rizzuto, R., and De Stefani, D. (2018). MICU3 is a tissue-specific enhancer of mitochondrial calcium uptake. Cell Death & Differentiation. 10.1038/s41418-018-0113-8.

82. Gottschalk, B., Koshenov, Z., Waldeck-Weiermair, M., Radulovic, S., Oflaz, F.E., Hird, M., Bachkoenig, O.A., Leitinger, G., Malli, R., and Graier, W.F. (2022). MICUl controls spatial membrane potential gradients and guides Ca2+ fluxes within mitochondrial substructures. Commun Biol 5,1-13. 10.1038/s42003-022-03606-3.

83. Dynes, J.L., Yeromin, A.V., and Cahalan, M.D. (2023). Photoswitching alters fluorescence readout ofjGCaMP8 Ca2+ indicators tethered to Orail channels. Proceedings of the National Academy of Sciences 120, e2309328120. 10.1073/pnas.2309328120.

84. Zhang, Y., R6zsa, M., Liang, Y., Bushey, D., Wei, Z., Zheng, J., Reep, D., Broussard, G.J., Tsang, A., Tsegaye, G., et al. (2023). Fast and sensitive GCaMP calcium indicators for imaging neural populations. Nature 615, 884–891. 10.1038/s41586-023-05828-9.

85. Nakai, J., Ohkura, M., and Imoto, K. (2001). A high signal-to-noise Ca2+ probe composed of a single green fluorescent protein. Nat Biotechnol 19, 137–141. 10.1038/84397.

86. Park, D., Veenstra, J.A., Park, J.H., and Taghert, P.H. (2008). Mapping Peptidergic Cells in Drosophila: Where DIMM Fits In. PLOS ONE 3, el 896. 10.1371/journal.pone.0001896.

87. Studier, F.W. (2005). Protein production by auto-induction in high-density shaking cultures. Protein Expression and Purification 41, 207–234. 10.1016/j.pep.2005.01.016.

88. Tsien, R., and Pozzan, T. (1989). Measurement of cytosolic free Ca2+ with quin2. Methods Enzymol 172, 230–262. 10.1016/s0076-6879(89)72017-6.

89. Pascual-Caro, C., and Juan-Sanz, J.de (2024). Monitoring of activity-driven trafficking of endogenous synaptic proteins through proximity labeling. PLOS Biology 22, e3002860. 10.1371/journal.pbio.3002860.

90. Vishwanath, A.A., Comyn, T., Chintaluri, C., Ramon-Duaso, C., Fan, R., Sivakumar, R., Lopez-Manzaneda, M., Preat, T., Vogels, T.P., Rangaraju, V., et al. (2024). Mitochondrial Ca2+ efilux controls neuronal metabolism and long-term memory across species. Preprint at bioRxiv, 10.1101/2024.02.01.578153 https://doi.org/10.1101/2024.02.01.578153.

91. Cuhadar, U., Calzado-Reyes, L., Pascual-Caro, C., Aberra, A.S., Ritzau-Jost, A., Aggarwal, A., lbata, K., Podgorski, K., Yuzaki, M., Geis, C., et al. (2024). Activity-driven synaptic translocation of LGil controls excitatory neurotransmission. Cell Reports 43. 10.1016/j.celrep.2024.114186.

92. Silva, B., Mantha, O.L., Schor, J., Pascual, A., Plaais, P.-Y., Pavlowsky, A., and Preat, T. (2022). Glia fuel neurons with locally synthesized ketone bodies to sustain memory under starvation. Nat Metab 4, 213–224. 10.1038/s42255-022-00528-6.

93. Hadjieconomou, D., King, G., Gaspar, P., Mineo, A., Blackie, L., Ameku, T., Studd, C., de Mendoza, A., Diao, F., White, B.H., et al. (2020). Enteric neurons increase maternal food intake during reproduction. Nature 587, 455–459. 10.1038/s41586-020-2866-8.

